# PBDE flame retardant exposure causes neurobehavioral and transcriptional effects in first-generation but not second-generation offspring fish

**DOI:** 10.1101/2025.04.28.651134

**Authors:** Nicole McNabb-Kelada, Tara Burke, Saro Jayaraman, Lesley Mills, Ashley De La Torre, Madison Francoeur, Hannah Schrader, Diane Nacci, Bryan Clark, Andrew Whitehead

## Abstract

Environmental pollutants can have lasting effects that extend beyond the exposure period, potentially impacting multiple generations. Despite the widespread presence of polybrominated diphenyl ether (PBDE) flame retardants in the environment including oceans globally, there is limited understanding of their potential for multigenerational toxicity. We investigated whether exposures to 2,2’,4,4’,5-pentaBDE (BDE-99), a predominant PBDE congener, can induce neurobehavioral and molecular effects across two generations using the marine vertebrate model fish *Fundulus heteroclitus* (Atlantic killifish). To assess how exposure route influences outcomes, we conducted two complementary experiments: in the progenitor exposure experiment, adult fish were exposed to BDE-99 through diet, leading to maternal transfer into F1 eggs, while in the direct embryonic exposure experiment, embryos were exposed to BDE-99 via waterborne exposure, achieving comparable embryonic doses. Both exposure routes resulted in altered photomotor responses in F1 larvae, but additional effects differed by exposure route. In the progenitor exposure, maternally exposed F1 juvenile fish exhibited reduced anxiety-like behavior and changes in brain gene expression, while in the direct embryonic exposure, F0 fish showed no detectable effects within their generation but caused behavioral alterations in F1 descendants. The lack of effects in direct F0 individuals suggests that neurobehavioral effects observed in maternally exposed F1 fish were not solely due to BDE-99 transfer but were influenced by additional maternal factors. In the progenitor exposure, neither behavioral nor gene expression effects persisted in the F2 generation. We conclude that maternal influences play a key role in shaping multigenerational effects of BDE-99 exposure, as indicated by the strong effects observed in maternally exposed F1 fish but not in F2 descendants, as well as the differences between maternally exposed fish and those exposed to comparable doses of BDE-99 alone.

## Introduction

Exposure to environmental stressors can leave a lasting imprint on organisms, with effects that extend beyond the direct exposure period, potentially impacting subsequent generations^1–3^. Factors such as poor nutrition, stress, and exposure to environmental toxicants can result in adverse health outcomes that are transmitted across generations^4–6^. This phenomenon, known as multi- and transgenerational effects, highlights the ability of organisms to pass on information about environmental perturbations to their offspring, potentially resulting in health impacts that emerge in later generations. Research has shown that various environmental contaminants that are commonly detected in marine environments including endocrine disruptors^7–10^, heavy metals^11–13^, and persistent organic pollutants^14–18^, can contribute to such heritable effects. However, despite increasing evidence of this phenomenon, much remains poorly understood, particularly the mechanisms underlying these multigenerational responses and the extent to which different classes of chemicals contribute to these outcomes. There is a critical need to incorporate multigenerational considerations into risk assessment frameworks to inform public health and environmental protection strategies.

Effects of toxicant exposure can propagate across generations through various mechanisms. Intergenerational effects (from parent to offspring) may result from altered maternal provisioning^19,20^, direct transfer of toxicants and metabolites^21,22^, changes in hormone and/or immune signaling^23–26^, and epigenetic modifications^27,28^. In oviparous fish, maternal provisioning influences offspring development by transferring both essential nutrients and harmful substances, including toxicants, to eggs, thereby exposing embryos at the earliest stages of life. Transgenerational inheritance, which is confirmed only when effects persist in an unexposed generation (F2 in oviparous fish), tend to not include contributions from maternal influences^5^ and rather implicate the influence of heritable epigenetic mechanisms such as changes in DNA methylation or histone modifications. Clues about contributing mechanisms emerge from examining effects across multiple generations: effects limited to F1 suggest maternal or direct exposure influences, whereas persistence in F2 suggests epigenetic inheritance^5,29^. To better understand whether and how exposure-induced impacts persist across generations, we assessed effects in both F1 and F2 generations.

Studies have shown that toxicants like vinclozolin^7^, dioxin^14^, DDT^15^, and PFAS^16^ can drive transgenerational inheritance, but there is still limited understanding of whether some major classes of persistent organic pollutants that are common in marine environments, such as polybrominated diphenyl ethers (PBDEs), induce similar effects. PBDEs are chemicals used as flame retardants in many consumer products, including plastics, electronics, furniture, textiles, and building materials. Since PBDEs are not chemically bonded to the materials to which they are added, they gradually leach into the environment^30^. PBDEs contaminate diverse environments, including marine environments globally, arriving in estuaries and oceans usually sorbed to particles through atmospheric deposition, river discharge, and directly through litter, including marine plastics, especially in coastal regions with large human presence^31–35^. Although penta-brominated diphenyl ethers (penta-BDEs) and octa-brominated diphenyl ethers (octa-BDEs) were largely phased out worldwide in the early 2000s due to toxicity concerns, deca-brominated diphenyl ether (deca-BDE) products are still legally manufactured in Asia^36,37^. Despite the phase-out, PBDEs are still being released into the environment through the disposal and recycling of electronic waste^38–41^. Additionally, these chemicals are highly lipophilic and capable of bioaccumulation and biomagnification in food chains, resulting in their persistence and ubiquitous presence in the environment, such as in air, house dust, freshwater and marine sediments, sewage sludge, soil, and biota including humans and wildlife^42–51^ including diverse marine species especially those consumed by humans^52–54^. PBDEs also exhibit significant capability for maternal transfer, being passed from pregnant females to offspring through the placenta and breast milk in mammals^55–58^, as well as through egg deposition in oviparous species like fish^59,60^, highlighting the potential risks to both human and wildlife species across multiple generations. In this study, we designed experiments to test whether exposure to 2,2’,4,4’,5-pentaBDE (BDE-99), a prevalent PBDE congener, results in perturbations in the offspring and grand-offspring of exposed fish.

Adverse health impacts of PBDEs have been extensively studied, particularly on neurological and endocrine systems. Among the PBDE congeners, BDE-99 is one of the most common congeners detected in salmon^61^ as well as in human samples, including serum and breast milk^62,63^. Studies have indicated adverse effects of BDE-99 exposure on development and neurobehavior across various species, including rats^64^, birds^65^, zebrafish^66^, and humans^67^. Endocrine-disrupting effects of BDE-99 exposure have also been reported, including altered thyroid hormone levels in rats^64^ and salmon^68^ and interactions with several hormone receptors in zebrafish, including PPARα, AhR2, PXR, glucocorticoid, and thyroid hormone receptors^69^. While there is substantial evidence highlighting the detrimental effects of PBDEs like BDE-99 in directly exposed individuals, research on their long-term multigenerational effects remains limited. Some studies have provided evidence of potential multigenerational toxicity, such as maternal transfer of PBDEs resulting in malformations in rats^70^ and decreased motor performance in zebrafish due to neurodevelopmental disruptions^71^. Also in zebrafish, exposure to a mixture of PCBs and PBDEs resulted in behavioral and molecular disruptions that persisted across multiple generations (F1-F4), but it remains unclear whether these transgenerational effects were driven by PCBs, PBDEs, or their combined action^72,73^. In birds, embryonic exposure to a PBDE mixture in American kestrels resulted in altered reproductive success and behaviors across generations^74^, while in zebra finches, *in ovo* exposure to BDE-99 had significant effects on the growth of second-generation offspring^75^. Given the persistence of PBDEs in the environment and limited data on their long-term effects across generations, we examined the multigenerational impacts of BDE-99 on neurobehavioral and molecular outcomes using Atlantic killifish (*Fundulus heteroclitus*).

The Atlantic killifish is ecologically important, nonmigratory, and prevalent in U.S. Atlantic coast salt marsh estuaries^76^, including areas contaminated with persistent pollutants^77^. Serving as both an ecological sentinel and a well-established model in ecotoxicology, *F. heteroclitus* has several attributes that make it suitable for studying multigenerational impacts of toxicant exposure. As an outbred, genetically diverse species, it may provide insights that are more representative of typical wild populations than those obtained from inbred, laboratory-adapted animal strains. Neurobiological mechanisms are highly conserved across vertebrates, making *F. heteroclitus* a relevant model for studying neurotoxicological effects in fish and tetrapods, including marine mammals and humans^78,79^.

To investigate the propagation of effects from BDE-99 exposure across two generations, we integrated brain transcriptomics with behavioral outcomes, given the known neurotoxicity of PBDEs. In a companion study we examined effects that persist from through to adulthood (McNabb-Kelada et al., unpublished), whereas in this study we focus on effects that propagate across generations. In this study we compared two distinct exposure regimes: maternal transfer and direct embryonic (waterborne) exposure. For maternal transfer, adult killifish (F0) were exposed to BDE-99 through diet for 64 days and spawned to produce the first generation (F1) of the progenitor exposure lineages (plus no-exposure control lineage). In parallel, we conducted 6-day waterborne exposures with embryos from unexposed parents to create the first generation (F0) of the direct embryonic exposure lineages (plus no-exposure control lineage). Embryos from both sets of lineages were raised in uncontaminated conditions and subsequently spawned to produce a second generation, allowing us to assess inheritance of effects. We evaluated a range of endpoints, including developmental morphology, hatch success, larval survival, photomotor response, anxiety-like behavior, and whole-brain genome-wide gene expression, across two generations and both exposure regimes. This approach was specifically designed to test two hypotheses: (1) that developmental exposure to BDE-99 causes intergenerational neurotoxicity, leading to altered behavior and gene expression patterns in the first generation, with some perturbations propagating to the second generation; and (2) that exposure route (maternal vs. environmental) influences these outcomes due to maternal factors in addition to direct toxicant transfer. We therefore predicted that outcomes between the two exposure routes would differ, reflecting the influence of these maternal effects. Since multigenerational impacts can lead to unpredictable phenotypic outcomes due to complex genetic, epigenetic, and environmental interactions, we measured genome-wide gene expression to identify potential disruptions in molecular functions and pathways, helping to generate hypotheses about alternative phenotypic effects that were not directly measured.

## Methods

We conducted two complementary experiments to investigate the multigenerational effects of exposure to BDE-99, using distinct exposure paradigms that differed in life stage and route of exposure (Figure 1). In Experiment 1, adult progenitor killifish were exposed to BDE-99 through diet, and effects were assessed across the next two generations. In Experiment 2, embryos were directly exposed to BDE-99 through waterborne exposure and achieved comparable doses as in Experiment 1, allowing for a comparison of whether maternal transfer vs. direct environmental exposure influences multigenerational effects. Though many of the methods were identical between experiments, we describe them sequentially below. More detailed methods are available in the supplemental materials (Supplemental Methods).

**Figure 1:**
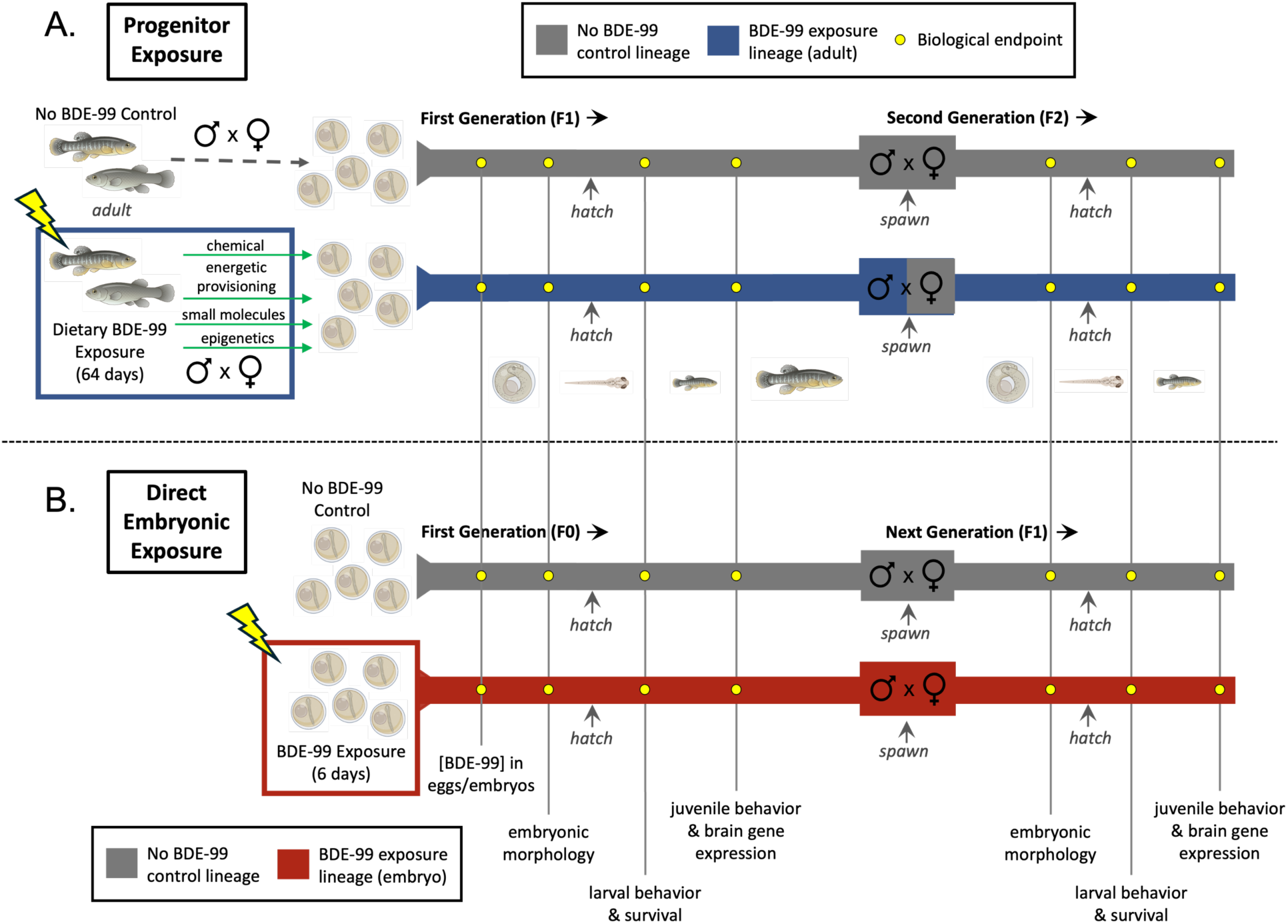
Multigenerational study paradigm of progenitor (A, Experiment 1) and direct embryonic (B, Experiment 2) exposure to BDE-99. Developing killifish embryos were exposed to BDE-99 either through A) transfer of the chemical from adult progenitors, or B) directly from their surrounding water. A) Adult progenitor (F0) fish were exposed to control or BDE-99 (blue box) diets for 64 days. First-generation (F1) descendants (gray line is control lineage, blue line is progenitor exposure lineage) were produced by mating males and females within each lineage and raised in uncontaminated conditions. To produce second-generation (F2) descendants, males in each lineage were mated with non-exposed females. B) For the direct exposure experiments, embryos from unexposed parents were exposed to control (gray line) or BDE-99 (red box) via waterborne exposure from 1-7 days post-fertilization (dpf). Those first-generation (F0) fish were raised in uncontaminated conditions until adulthood (gray line is control lineage, red line is direct embryonic exposure lineage), and then males and females within each lineage were mated to produce next-generation (F1) descendants. Biological endpoints were measured across two generations in both exposure experiments at the same time points through development. Fish and embryo illustrations sourced from BioRender.com.

### Progenitor Exposure (Experiment 1) Chronological Overview

Adult wild-caught killifish were exposed to either control or BDE-99-contaminated diets for 64 days during the breeding season. Throughout this exposure period, the exposed (F0) progenitor fish were spawned to produce the parentally exposed first (F1) generation. The F1 fish were reared under uncontaminated conditions until adulthood (2 years) and subsequently spawned to produce the second (F2) generation (Figure 1A). A suite of endpoints was assessed across both generations: developmental abnormalities at 10 days post-fertilization (dpf), hatch success at 14 dpf, larval survival until 14 days post-hatch (dph), larval photomotor response (light/dark transition) at 14 dph, and juvenile anxiety-like behavior (novel tank diving) at 4-6 months post-hatch. After behavioral testing was completed, brains from F1 and F2 fish were flash-frozen and preserved for transcriptomics.

### Adult Fish Collection and Husbandry

Adult Atlantic killifish (∼10 g wet weight (ww)) were collected from Jerusalem, Narragansett, RI, USA (41°22’58.4” N, 71°31’09.6” W)^80^ and housed at the US Environmental Protection Agency (EPA) laboratory in Narragansett in tanks (300 L, 80 gal) with continuous flow of 23°C, 5 µm-filtered, clean seawater, under a 14:10 h light:dark cycle and fed TetraMin Tropical Flakes fish food (Tetra, Blacksburg, VA, USA) *ad libitum* until their use in this study one month later. Before the experiment, fish were redistributed into 38-L (10 gal) tanks. Each treatment consisted of 3 replicate tanks, with each tank containing 14 fish (8 females and 6 males). All procedures were in accordance with EPA-approved Animal Care and Use Protocols (IACUC, ACUP # Eco23-03-002, Eco23-11-001, and Eco23-07-001).

### Diet Preparation for Progenitor BDE-99 Exposure

A BDE-99 (2,2’,4,4’,5-pentabromodiphenyl ether; >99% pure) stock solution (2.5 mg/mL) was prepared. To prepare the BDE-99_MED and BDE-99_HIGH dosing solutions, 52 µL or 209 µL of the BDE-99 stock was added to 1348 µL or 1191 µL of acetone, respectively (final volumes of 1.4 mL). Diet consisted of ground TetraMin Tropical Flakes (40 g) amended with either carrier (acetone, 1.4 mL) or BDE-99 stock dissolved in acetone (1.4 mL). A nutritional mixture (including blended spinach, brine shrimp, krill, mackerel, sardines) supplemented the experimental diet, which was finally solidified with gelatin.

### Progenitor Dietary Exposure to BDE-99

Fish were fed control or contaminated diets (∼15% of body weight) 5 days per week and control diet the other 2 days a week for 64 days. Dietary treatments consisted of control and 2 concentrations of BDE-99 (37.5 and 150 ng/g fish ww/day). Target internal doses of BDE-99 (verified as described below) were 1 and 4 μg/g fish dry weight (dw).

### Exposed Generation Endpoints

Standard length (SL, mm) and whole fish wet weight (ww, g) were recorded for individual fish at the start (day 0) and end (day 65) of the exposure, after which Fulton’s condition index was calculated (ww/SL^3^). During this period, F0 fish were spawned at multiple time points to generate the F1 generation. After 64 days of feeding, all F0 (progenitor) fish were euthanized, after which wet weights of gonads, liver, and abdominal fat, were measured and used to calculate the health indices: gonadosomatic index (GSI), hepatosomatic index (HSI), and abdominal fat somatic index (AFSI). Carcasses were then vacuum-sealed, flash-frozen, and stored at -80°C for BDE-99 body burden quantification. Exposure effects on growth and health indices were tested using two-way Analysis of Variance (ANOVA) with tank as the unit of replication, where main effects included BDE-99 concentration and sex, and their interaction. Before ANOVA, data normality was tested (Shapiro-Wilk test). Sex effects on body burden were tested (Welch’s t-tests). Statistical analyses were conducted using GraphPad Prism version 10.4.1 for macOS (GraphPad Software, Boston, MA, USA).

### Spawning

Fish were manually strip-spawned 5 times following their semi-lunar spawning cycle^81^ through the dietary exposure period to record reproductive success and egg counts. Fertilization to create the F1 generation included eggs and milt from fish from a single tank and kept separate so that subsequent matings (to form the F2) did not include close relatives. Embryos were then incubated at 23°C (14:10 h light:dark cycle). Fertilization success was recorded at 4 dpf, and unfertilized eggs were archived for chemical analysis. Exposure effects on fecundity were analyzed using a two-way repeated measures ANOVA with tank as the unit of replication. Fertilization success (percentage of fertilized F1 eggs per tank) was analyzed using a mixed-effects model with Greenhouse-Geisser correction (to account for possible sphericity violations). Statistical models included main effects of BDE-99 concentration, experimental day, and their interaction. Two-year-old F1 fish were spawned to produce the F2 generation. Due to low female numbers in F1 exposure and control lineages, males were crossed with external control females. As a result, only paternally-mediated intergenerational effects were assessed in F2-generation fish. Eggs collected from 20-36 females were pooled, divided into approximately equal groups, and fertilized with pooled milt from males within each BDE-99 exposure lineage. Fertilization measurements and rearing conditions were the same as in F1.

### Chemical Analysis

Accelerated Solvent Extraction was used to extract BDE-99 from freeze-dried F0 fish carcass samples and treatment diets^82^. Approximately 0.2 g of carcass was ground with a mortar and pestle, spiked with a known amount of IS, and extracted with acetone:hexane (50:50). A small portion (10%) was transferred to a pre-weighed aluminum weighing pan for lipid determination, and the remaining extract was treated with concentrated sulfuric acid. F1 fish eggs from progenitor exposure (Experiment 1) and F0 embryos from direct embryonic exposure (Experiment 2) (40-150 mg ww) were similarly treated.

BDE-99 concentrations in adult and early-life samples were measured by Gas Chromatography-Mass Spectrometry (GC-MS) using an Agilent Technologies (Santa Clara, CA, USA) 6890N GC with a 5973 mass selective detector and an Agilent J&W DB-5ms Ultra Inert analytical column (15 m x 0.25 mm x 0.25 µm). Calibration standard solutions of BDE-99 (50-2500 ng/µL) were prepared. ChemStation software was used for data processing. Procedural blanks showed no contaminants. Analytical duplicates had a relative standard deviation (RSD) of <7%, and matrix spike recoveries ranged from 90% to 104%.

### Embryo-Larval Rearing and Early-Life Endpoints

Embryo-Larval Assays (ELA) evaluated the developmental and survival effects of BDE-99 exposure lineage^83,84^. Embryos were housed in clean seawater refreshed daily. At 10 dpf (late organogenesis, stage 34^85^), embryos were screened (presence/absence) for developmental abnormalities, including pericardial edema, heart abnormalities, hemorrhaging in the head or tail, and reduced body or head size^86^. Treatment effects on developmental abnormalities were analyzed using the Kruskal-Wallis test. Treatment effects on hatching success and larval survival (until 14 dph) were analyzed using Fisher’s exact tests.

### Larval/Juvenile Behavioral Endpoints

Light/dark transition assays were conducted at 14 dph to assess larval photomotor responses. Plates containing larvae were placed on an infrared backlight inside a custom-built black box, and movement was recorded from above (Basler acA1300-60gmNIR camera). EthoVision XT software was used to track activity. Assays began with 30-min light habituation (100% illumination), followed by alternating light and dark phases: a 10-min initial dark phase (0% illumination), then two cycles of alternating 10-min phases in light (100% illumination) and dark. *Total distance moved* was calculated as cm per 10-min interval, with the initial habituation phase excluded. Following Greenhouse-Geisser correction, data were analyzed using two-way repeated measures ANOVA with individual larval fish as the subject (biological replicate), where main effects were BDE-99 exposure lineage, light phase, and their interaction, followed by Dunnett’s post-hoc tests.

Novel tank diving assays were performed to evaluate anxiety-like behaviors in juvenile fish (4-6 months post-hatch)^87^. The experiment used two adjacent rectangular tanks, each filled with ∼4.5 L seawater (21.5-23°C). Behavioral trials were recorded (10-min sessions) using a camera placed 120 cm from the front of the tanks, with tracking and behavior analyzed by EthoVision XT software. Behavioral measurements included *total distance moved* (cm per 10 min), *duration spent in the top zone* (percent time), *latency to enter the top zone* (seconds), and *number of transitions to the top zone*. Subsets of fish were then euthanized (MS-222) and brain tissues preserved for transcriptomics analysis. Data were tested for normality (Shapiro-Wilk test) and then analyzed using one-way ANOVA, followed by Tukey’s post-hoc tests. Endpoints that failed normality were analyzed using the Kruskal-Wallis test, followed by Dunn’s multiple comparisons test.

### Transcriptomics

Juvenile brains were homogenized (Omni Bead Ruptor) and mRNA extracted using the Dynabeads^TM^ mRNA DIRECT^TM^ Purification Kit (Invitrogen, Waltham, MA, USA). Strand-specific RNA-seq libraries were prepared using the Breath Adapter Directional sequencing (BrAD-seq) method^88^. Individual libraries were uniquely indexed, pooled, and sequenced across six lanes of Illumina HiSeq 4000 (PE-150) (UC Davis Genome Center). Each sample yielded ∼9 million raw reads. F1 samples included 4-5 biological replicates per treatment (except n=3 for the 0.85 µg/g group), for a total of 27 samples. F2 samples included 6 replicates per treatment (except n=2 for the 3.50 µg/g group), for a total of 32 samples. Six exposure lineages were represented in each generation: 0, 0.85, 1.20, 2.00, and 3.50 µg/g in F1 and F2, with 0.57 and 0.47 µg/g in F1 and F2, respectively. Raw reads were quality checked (FastQC^89^), trimmed (fastp^90^), and mapped to the *F. heteroclitus* reference genome (GenBank MU-UCD_Fhet_4.1)^91^ using Salmon^92^.

We applied weighted gene correlation network analysis (WGCNA)^93^ to identify modules of co-expressed genes that varied with experimental endpoints, conducted separately for the F1 and F2 generations. Raw counts were normalized using variance-stabilizing transformation in DESeq2^94^. The top 10% most variable genes were selected (3,164 per dataset) as recommended by WGCNA. Gene modules were identified using hierarchical clustering and dynamic tree cutting. For each module, eigengene values (PC1 of module expression) were extracted and tested for statistical association with treatment (BDE-99 exposure lineage) using one-way ANOVA (FDR-adjusted p<0.05). Modules with significant treatment effects were tested for Gene Ontology (GO) enrichment using a Mann-Whitney *U* test^95^. All analyses were conducted in R (version 4.4.2)^96^ and RStudio (version 2024.09.1+394).

### Direct Embryonic Exposure (Experiment 2) Chronological Overview

Fish embryos collected from unexposed parents were subjected to 6-day waterborne exposures to control or BDE-99 from 1-7 dpf (Figure 1B). These directly exposed F0 embryos were then reared to adulthood under uncontaminated conditions and subsequently spawned to produce the next (F1) generation. The same suite of developmental and behavioral endpoints was measured following these direct embryonic exposures as in the direct progenitor exposures (Experiment 1) described above: developmental abnormalities at 10 dpf, hatch success at 14 dpf, larval survival until 14 dph, larval photomotor response (light/dark transition) at 14 dph, and juvenile anxiety-like behavior (novel tank diving) at 4-6 months post-hatch. Following behavioral testing, whole brains from the F1 generation were collected and archived for RNA-seq.

### Embryo Collection

Embryos were from mass-spawned adults collected from Scorton Creek, Sandwich, MA (41°45’53.6” N, 70°28’48.0” W)^97–100^ that had been housed at the US EPA facility. Four separate breeding tanks were used to produce four groups of embryos, which were kept separate to prevent mating between close relatives to produce the next (F1) generation. From each breeding tank, eggs were collected from 20-30 females, pooled, and fertilized with milt pooled from 20-30 males from the same tank.

### Direct Embryonic Exposure to BDE-99

A BDE-99 stock solution (2.5 mg/mL) was prepared as described above. Fertilized embryos from each of the four breeding groups were equally divided into three treatments (control and two BDE-99 concentrations) and then into two replicates per treatment. Each exposure experiment consisted of 24 jars (4 breeding groups x 3 treatments x 2 replicates per treatment). Embryos were mass exposed from 1 to 7 dpf in glass jars at a density of 1 embryo per 2 mL of seawater, amended with 1% dosing solution containing either carrier (acetone, control) or BDE-99 in acetone. BDE-99 concentrations were 6.2 and 26 ng/mL (labeled MED and HI, respectively). The entire exposure experiment was repeated across three separate spawning events, following the semi-lunar spawning cycle. Individuals from one of the four breeding groups and one of the three spawning cycles were archived for BDE-99 body burden analysis, such that embryo dosing could be compared between experiments 1 and 2.

### Larval/Juvenile and Behavioral Endpoints

The same endpoints were collected for these direct embryonic exposures as for the progenitor exposures (Experiment 1) as described above, including developmental abnormality, hatch success, larval photomotor response, larval survival, and juvenile anxiety-like behavior. Statistical analyses were also identical to those described above for Experiment 1. Data from the F0 and F1 generations were analyzed separately.

### Rearing to Adult and Spawning

Fish from the direct embryonic exposure study were reared as described above for the direct progenitor exposures (Experiment 1). The F1 generation was produced by crossing males and females from the different replicate groups within each exposure lineage. Therefore, effects in the F1 generation were both paternally and maternally mediated. Eggs were collected via manual strip-spawning, pooled within each replicate group, and fertilized with milt pooled from males from another replicate group in the same exposure lineage.

### Transcriptomics

Whole brain tissues from direct embryonic exposure F1 juvenile fish were processed, and RNA-seq data collected and analyzed (WGCNA), as described above for Experiment 1. Each sample yielded approximately 9.5 million raw reads. There were 6 replicates each from 3 breeding groups per exposure lineage, except for the MED BDE-99 lineage, which had 3 and 4 replicates in 2 of the breeding groups, for a total of 49 samples.

## Results

### Progenitor Exposure (Experiment 1): F0 Growth, Health, Fecundity, Fertility

Exposure to BDE-99 had no significant impact on growth or health indices in diet-exposed adults (Figure S1). Tissue indices (GSI, HSI, and AFSI) were significantly higher in females than males (p=0.0001, p<0.0001, and p=0.0431, respectively). Fecundity and fertility were variable over time, but BDE-99 had no significant effect on fecundity or fertilization success of diet-exposed adult progenitor fish (Figure S2).

### Progenitor Exposure: F0 BDE-99 Body Burdens and Diet Concentrations

Bioaccumulation of BDE-99 through diet in F0 adults was confirmed (Figure S3). Measured concentrations in males and females in the CTL, MED, and HI groups were 0.0106 (± 0.0062) µg/g and 0.0133 (± 0.0072) µg/g dw, 1.17 (± 0.20) µg/g dw and 0.933 (± 0.058) µg/g dw, and 4.47 (± 0.14) µg/g dw and 3.80 (± 0.32) µg/g dw, respectively, which were close to dose targets (1 and 4 µg/g dw). Male and female body burdens were not different. Measured concentrations of BDE-99 in treatment diets were not detected (ND), 0.895 µg/g dw, and 3.99 µg/g dw in the CTL, MED, and HI, respectively.

### Maternal Transfer of BDE-99 to F1

BDE-99 was detected in unfertilized F1 eggs from the progenitor exposure, confirming maternal transfer (Figure 2A). F1 doses increased with duration of maternal exposure and dietary concentration, consistent with bioaccumulation in progenitors and subsequent maternal transfer into eggs. F1 eggs from lineages for which we subsequently collected data on hatching, larval survival, and photomotor response had measured BDE-99 concentrations of 0 (control), 0.42, 0.47, 0.57, and 2.40 µg/g dw. F1 eggs from lineages for which we subsequently collected neurophysiological data (juvenile novel tank testing, brain gene expression) had concentrations of 0 (control), 0.57, 0.85, 1.20, 2.00, and 3.50 µg/g dw. F1 eggs from lineages used for F2 production had concentrations of 0 (control), 0.47, 0.85, 1.20, 2.00, and 3.50 µg/g dw.

**Figure 2:**
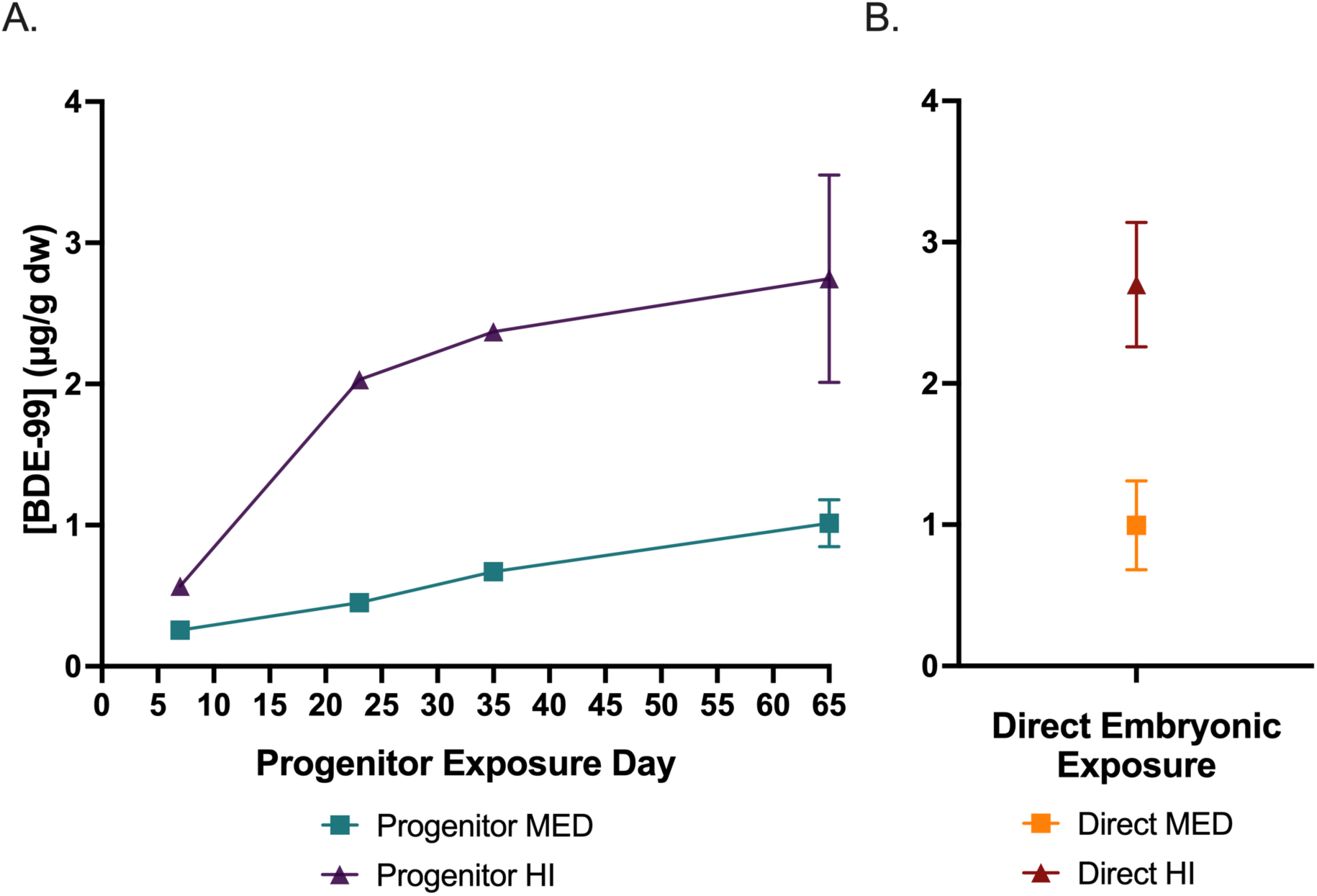
Maternal transfer of BDE-99 to F1 eggs from progenitor exposure (A) compared with direct waterborne exposure of F0 embryos (B). A) BDE-99 doses (µg/g dry weight (dw)) in unfertilized F1 eggs produced by progenitor (F0) fish over the 64-day dietary exposure period to BDE-99. Symbol colors and shapes indicate treatment groups: Progenitor BDE-99 MED (teal squares), Progenitor BDE-99 HI (purple triangles). B) BDE-99 doses (µg/g dw) in F0 embryos following direct waterborne exposure from 1-7 days post-fertilization (dpf) (mean ± SE). Symbol colors and shapes indicate treatment groups: Direct BDE-99 MED (orange square) and Direct BDE-99 HI (red triangle).

In the direct embryonic exposure experiment, BDE-99 was measured in subsets of F0 embryos following waterborne exposure from 1-7 dpf (Figure 2B). The mean body burdens achieved through direct environmental (waterborne) exposure (1.00 µg/g and 2.70 µg/g) (Experiment 2) successfully approximated the mean BDE-99 doses achieved via maternal transfer by the end of the dietary exposure period (1.01 µg/g and 2.75 µg/g) (Experiment 1).

### Early-Life Development and Survival Following BDE-99 Exposure

Progenitor dietary exposure to BDE-99 caused no effects on offspring developmental abnormalities, hatching, or larval survival, in either the F1 or F2 generations (Table S1), compared to controls. Similarly, direct embryonic (waterborne) exposure to BDE-99 had no significant effects on the same endpoints.

### Progenitor Exposure: Photomotor Responses in F1 and F2 Descendants

BDE-99 exposure lineage, defined as the progenitor exposure history inherited by each generation, significantly affected larval photomotor responses in the F1 generation, while effects in the F2 generation were minimal. In the F1 generation, BDE-99 exposure lineage had a significant effect on locomotor activity (p<0.0001), with a trend toward an interaction between exposure lineage and illumination phase (p=0.0524) (Figure 3A). Post-hoc comparisons revealed significant hypoactivity in F1 larvae from the 0.42, 0.47, and 0.57 µg/g exposure lineages in most illumination phases (3 out of 5), while larvae from the highest exposure lineage (2.40 µg/g) showed no significant differences from controls.

**Figure 3:**
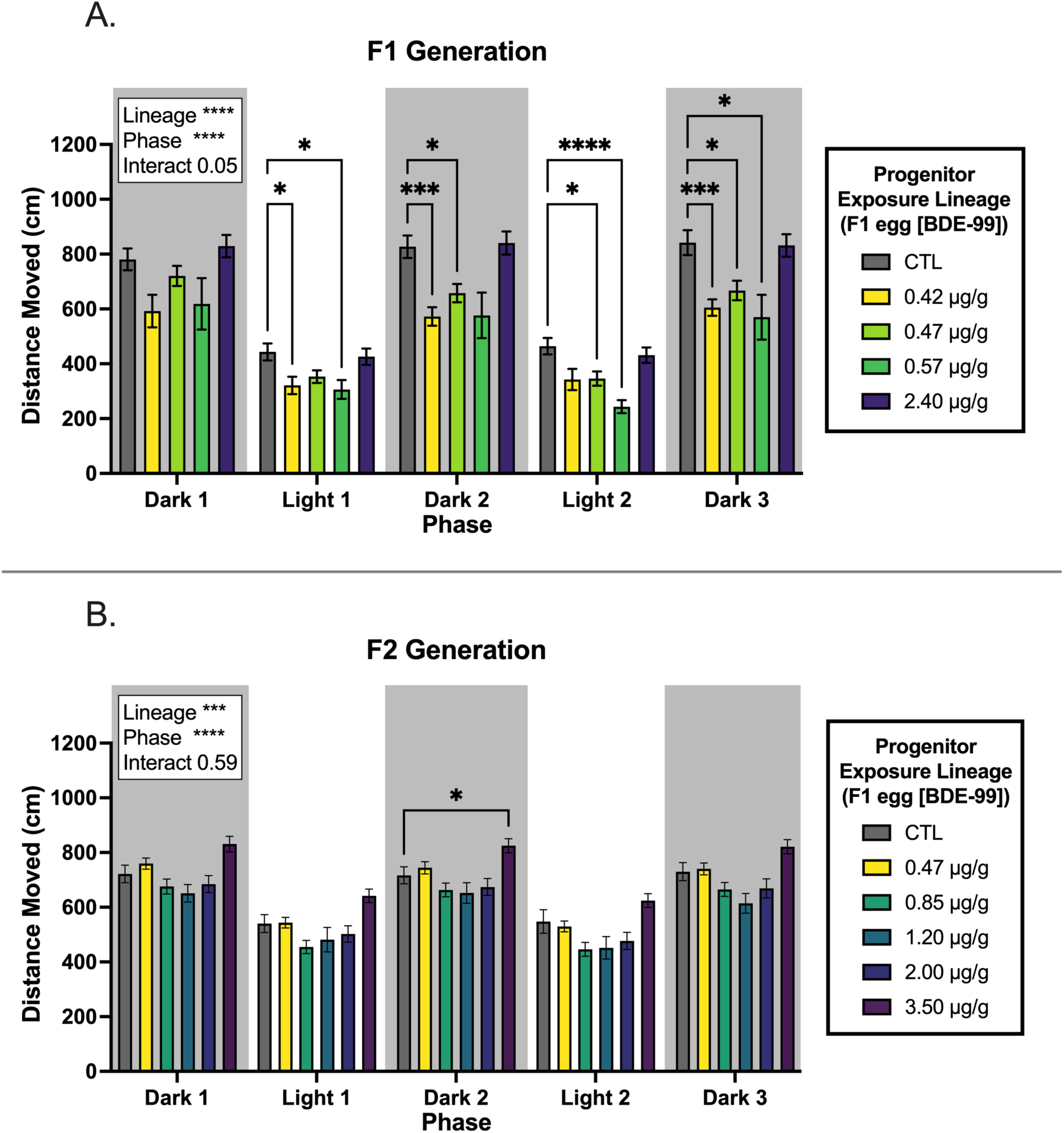
Larval photomotor response in F1 and F2 descendants of progenitor (F0) fish exposed to BDE-99 through diet. Total distance moved (cm) by F1 (A) and F2 (B) larvae during alternating light and dark phases. A) F1 larvae originated from control (CTL; n=23) or exposure lineages with the following BDE-99 concentrations (µg/g dry weight (dw)) measured in eggs: 0.42 (n=8), 0.47 (n=18), 0.57 (n=13), and 2.40 (n=17). B) F2 larvae originated from control (CTL; n=18) or exposure lineages with the following BDE-99 concentrations (µg/g dw) measured in F1 eggs: 0.47 (n=46), 0.85 (n=45), 1.20 (n=21), 2.00 (n=32), 3.50 (n=20). Within a plot, outcomes (p-values) for statistical tests are shown for the two main effects (progenitor exposure lineage and illumination phase) and their interaction. P<0.05, p<0.001, and p<0.0001 are indicated by *, ***, and ****, respectively. Brackets with asterisks indicate significant differences from CTL following post-hoc tests. Error bars represent standard error of the mean (SEM).

In contrast, the F2 generation exhibited more subtle effects caused by progenitor exposure (Figure 3B). There was a significant main effect of BDE-99 exposure lineage (p=0.0002), but no interaction with illumination phase (p=0.5883). The only significant lineage effect from post-hoc tests was increased movement (hyperactivity) in Dark 2 for larvae from the 3.50 µg/g BDE-99 lineage compared to controls (p=0.0391).

### Progenitor Exposure: Anxiety-Like Behavior Effects in F1 and F2 Descendants

Progenitor exposure to BDE-99 perturbed multiple dimensions of juvenile behavior in F1 descendants but not in F2 descendants. In the F1 generation, BDE-99 exposure lineage had a significant effect on *total distance moved* (p=0.0130), with hyperactivity observed in fish from the 0.85 and 1.20 µg/g lineages when compared to controls (p=0.0155 and 0.0184, respectively) (Figure 4A). F1 fish from multiple exposure lineages exhibited behavioral patterns consistent with reduced anxiety-like behavior, including increased *duration spent in the top zone* (p=0.0001) and more frequent *transitions to the top zone* (p=0.0011). Post-hoc comparisons revealed that fish from the 0.57, 0.85, 1.20, and 2.00 µg/g lineages spent more time in the top zone relative to controls (p=0.0010, 0.0275, 0.0057, and 0.0014, respectively) (Figure 4B). Similarly, fish from the same exposure lineages made more transitions to the top zone than controls (p=0.0086, 0.0091, 0.0064, and 0.0095, respectively) (Figure 4C). *Latency to enter the top zone* was also affected by exposure lineage (p=0.0001) (Figure 4D). Fish from the 0.57 and 0.85 µg/g lineages entered the top zone faster than controls (p=0.0036 and 0.0061, respectively). Additionally, latency was lower in the 0.57 and 0.85 µ/g lineages compared to 3.50 µ/g (p=0.0118 and 0.0218, respectively), suggesting a possible non-monotonic dose-response effect.

**Figure 4:**
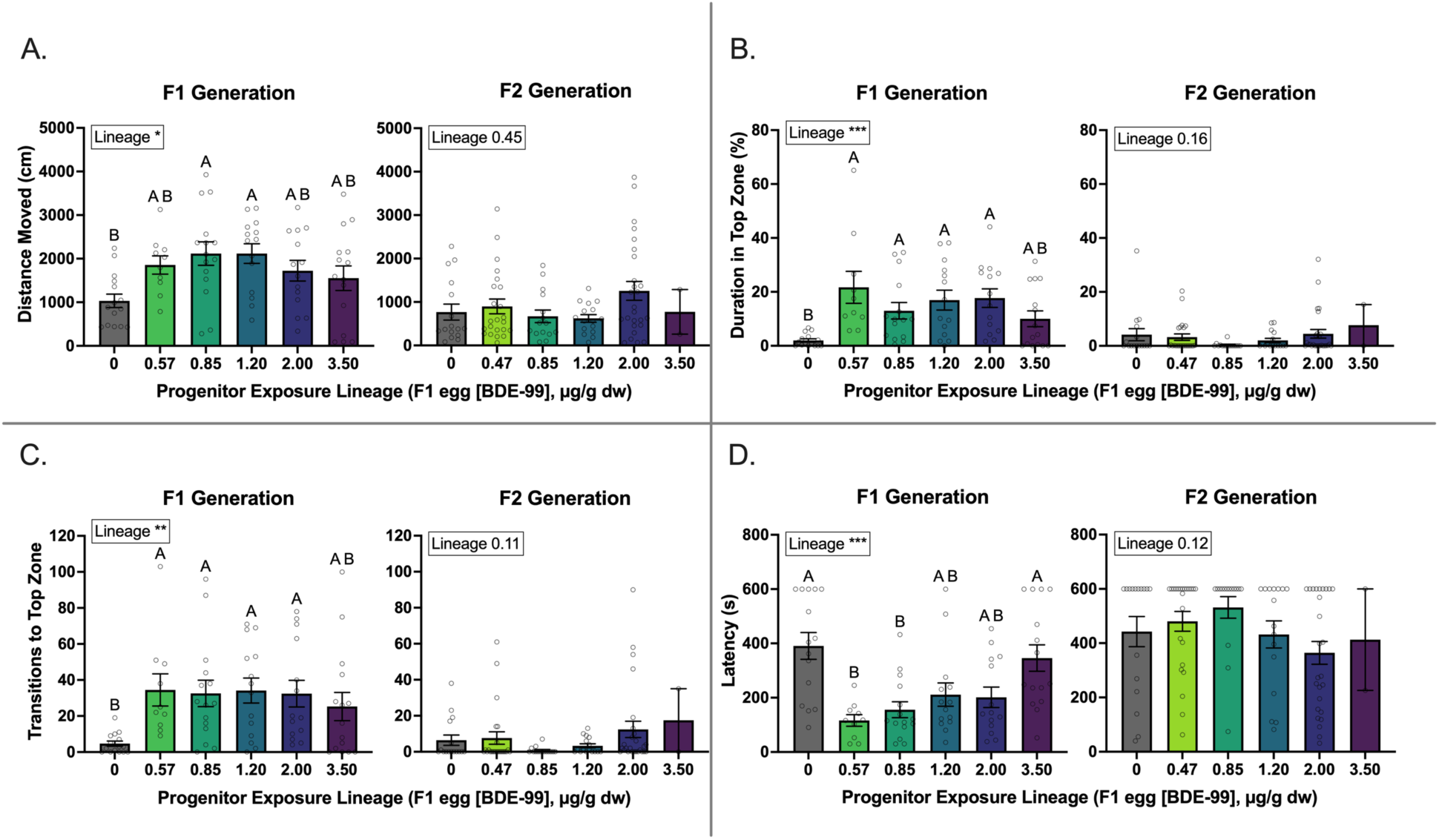
Juvenile anxiety-like behavior in F1 and F2 descendants of progenitor (F0) fish exposed to BDE-99 through diet. Variables included total distance moved (cm) (A), duration (% of time) spent in the top zone of the tank (B), number of transitions to the top zone (C), and latency (seconds) to enter the top zone (D). Fish that never entered the top zone were assigned the maximum latency of 600 seconds (duration of the assay). F1 fish originated from progenitor exposure lineages with the following BDE-99 concentrations (µg/g dry weight (dw)) measured in eggs: 0 (control; n=15), 0.57 (n=10), 0.85 (n=15), 1.20 (n=14), 2.00 (n=14), and 3.50 (n=15). F2 fish originated from lineages with the following BDE-99 concentrations (µg/g dw) measured in F1 eggs: 0 (control; n=16), 0.47 (n=23), 0.85 (n=15), 1.20 (n=16), 2.00 (n=26), and 3.50 (n=2). Within a plot, the statistical test outcome (p-value) is shown for the main effect of exposure lineage. P<0.05, p<0.01, and p<0.001 are indicated by *, **, and ***, respectively. The same letter above each bar indicates no significant difference between exposure lineage means following post-hoc tests. Error bars represent standard error of the mean (SEM).

Across behavioral variables, the 0.85 µg/g lineage showed the strongest effects, with 4 out of 4 variables significantly different from controls. The 0.57 and 1.20 µg/g lineages exhibited significant differences in 3 out of 4 variables, while the 2.00 µg/g group showed effects in 2 out of 4 variables. In contrast, though the highest dose lineage (3.50 µg/g) showed similar trends to lower dose lineages, there were no statistically significant behavioral differences between this lineage and controls (0 out of 4 variables). In contrast, no significant effects of BDE-99 exposure lineage were detected in the F2 generation. We conclude that progenitor exposure to BDE-99 affects juvenile behavior in the next generation, but these effects do not propagate into a second generation.

### Progenitor Exposure: Brain Transcriptome Responses in F1 and F2 Descendants

Progenitor exposures caused changes in brain gene expression in F1 descendant juveniles, but not F2 descendants. From F1 RNA-seq data, we detected nine co-expressed gene modules, each containing between 58 and 844 genes. Four modules were significantly associated with BDE-99 exposure lineage (adjusted p<0.05), with two modules containing functionally enriched gene sets.

The largest (turquoise) module (844 genes) had the strongest association (lowest p-value) with BDE-99 exposure lineage (adjusted p<0.0001). Principal Component Analysis (PCA) revealed that all three samples from the 0.85 µg/g lineage were transcriptionally distinct (Figure 5A). Transcription in samples from the 3.50 µg/g lineage also diverged from other lineages. Functional enrichment analysis indicated that genes in this module were primarily associated with immune system processes (e.g., activation of immune response, leukocyte activation, and cytokine-mediated signaling), and pathways related to cell communication and intracellular signaling (Figure 5B).

**Figure 5:**
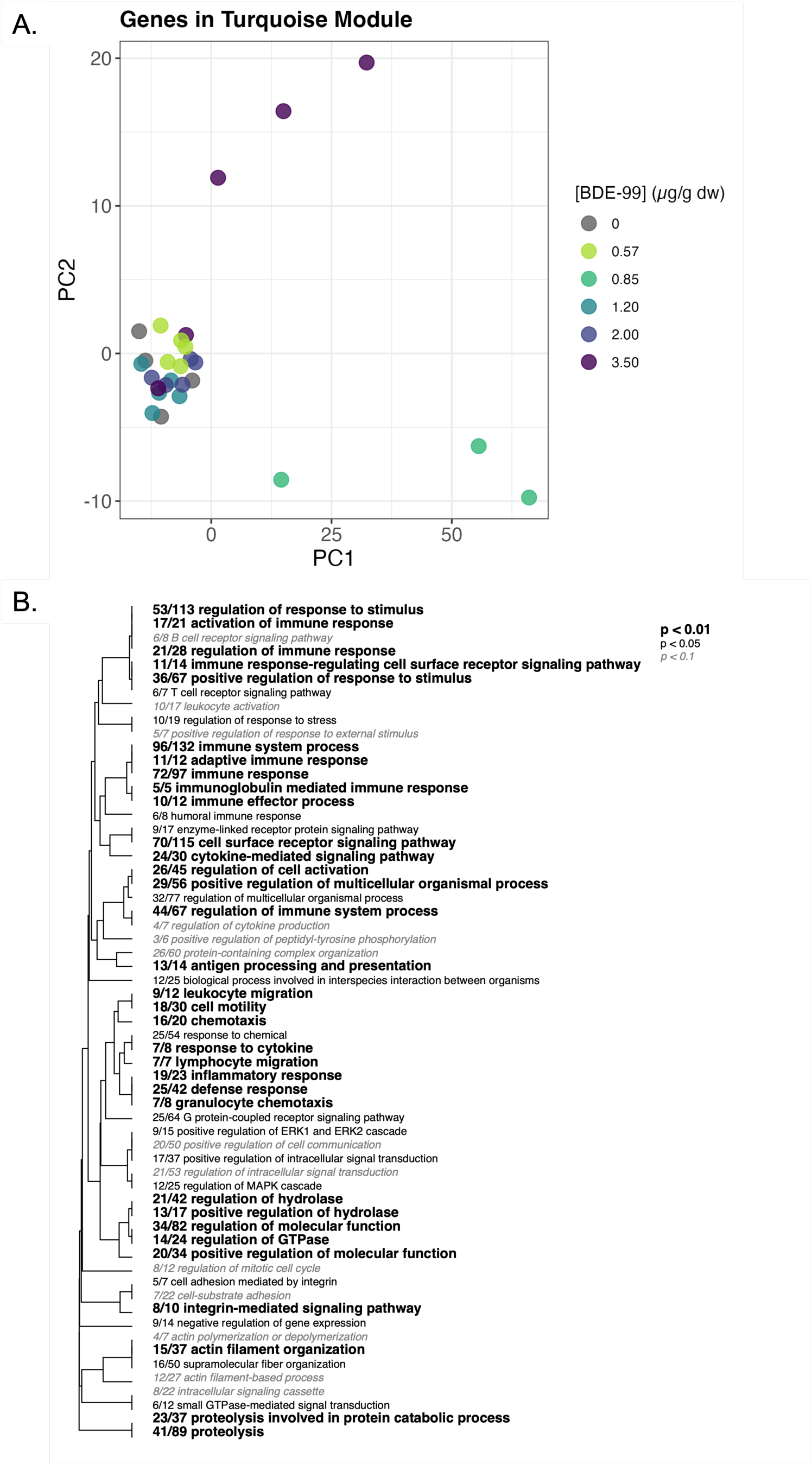
Principal Component Analysis (PCA) of whole-brain gene expression in the Turquoise module and associated Gene Ontology (GO) enrichment analysis in juvenile F1 fish developmentally exposed to BDE-99 via maternal transfer. A) PCA represents the variance in the expression of genes (n=844) assigned to the Turquoise module across samples. Points represent individual samples, colored by BDE-99 exposure lineage, including 0 (control; n=4), 0.57 (n=5), 0.85 (n=3), 1.20 (n=5), 2.00 (n=5), and 3.50 µ/g (n=5). B) Gene Ontology (biological processes) enrichment for the Turquoise module. The dendrogram represents hierarchical clustering of enriched GO terms based on shared genes. Fractions indicate the number of genes exceeding the unadjusted p<0.05 threshold relative to the total number of genes assigned to that category. GO terms are visually distinguished by statistical significance, with large, bolded text for p<0.01, regular text for p<0.05, and small, italicized text for p<0.1. GO terms are ordered by clustering similarity.

The Red module (146 genes) was also significantly associated with BDE-99 exposure lineage (p<0.0001). PCA showed distinct clustering of samples from the 0.57 and 0.85 µg/g BDE-99 exposure lineages, with the 0.57 µg/g group showing the most pronounced separation (Figure 6A). Genes in this module were enriched for processes related to transcriptional regulation, circadian rhythm, and metabolism, and for negative regulation of biosynthetic processes and response to environmental stimuli (Figure 6B).

**Figure 6:**
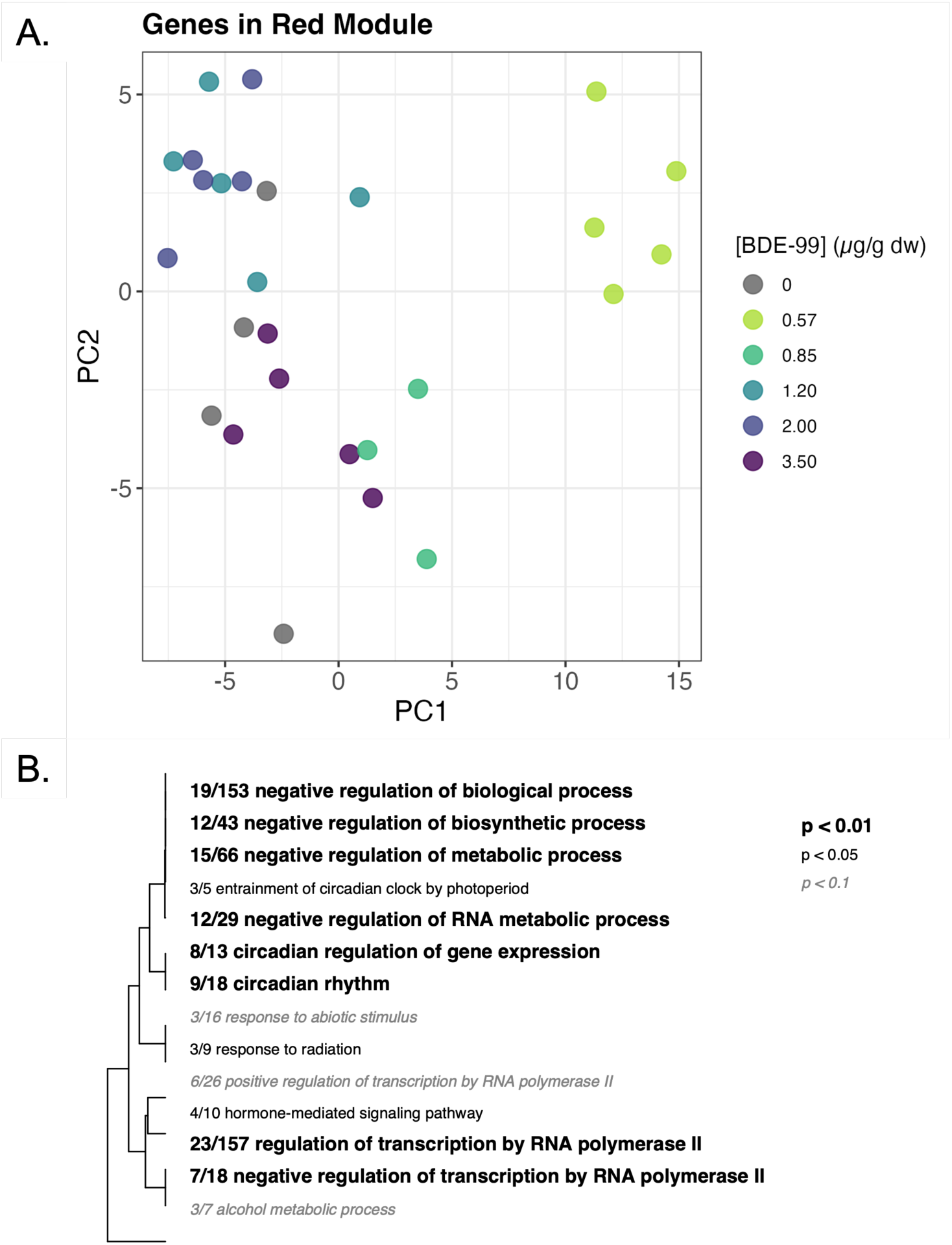
Principal Component Analysis (PCA) of whole-brain gene expression in the Red module and associated Gene Ontology (GO) enrichment analysis in juvenile F1 fish developmentally exposed to BDE-99 via maternal transfer. A) PCA represents the variance in the expression of genes (n=146) assigned to the Red module across samples. Points represent individual samples, colored by BDE-99 exposure lineage, including 0 (control; n=4), 0.57 (n=5), 0.85 (n=3), 1.20 (n=5), 2.00 (n=5), and 3.50 µ/g (n=5). B) Gene Ontology (biological processes) enrichment for the Red module. The dendrogram represents hierarchical clustering of enriched GO terms based on shared genes. Fractions indicate the number of genes exceeding the unadjusted p<0.05 threshold relative to the total number of genes assigned to that category. GO terms are visually distinguished by statistical significance, with large, bolded text for p<0.01, regular text for p<0.05, and small, italicized text for p<0.1. GO terms are ordered by clustering similarity.

From the F2 generation data, we detected 15 co-expressed gene modules containing 55 to 361 genes each, but none of these modules showed significant associations with BDE-99 exposure lineage (adjusted p>0.05). This pattern is consistent with behavioral data showing lineage effects in F1 offspring but not in the F2 generation, supporting the conclusion that adult exposures to BDE-99 have multigenerational impacts on neurobehavioral processes that span one generation but not two.

### Direct Embryonic Exposure (Experiment 2): Photomotor Responses in F0 and Their F1 Descendants

Direct embryonic exposure had a subtle but non-significant effect on photomotor function in larvae but had significant impacts on photomotor responses in F1 descendants. In the F0 (directly exposed) generation, there was a subtle but not statistically significant main effect of direct embryonic BDE-99 exposure (p=0.0723) (Figure 7A), trending toward hyperactivity. In contrast, direct embryonic BDE-99 exposure caused pronounced hypoactivity in F1 descendants (main effect of exposure lineage p<0.0001, interaction between exposure lineage and illumination phase p<0.0001) (Figure 7B). Post-hoc comparisons revealed that larvae from the BDE-99 HI (2.7 µg/g) lineage exhibited significant hypoactivity compared to control during both light and dark phases, and larvae from the BDE-99 MED (1.0 µg/g) lineage exhibited hypoactivity during light phases. Hypoactivity was greater in the HI lineage than in the MED lineage.

**Figure 7:**
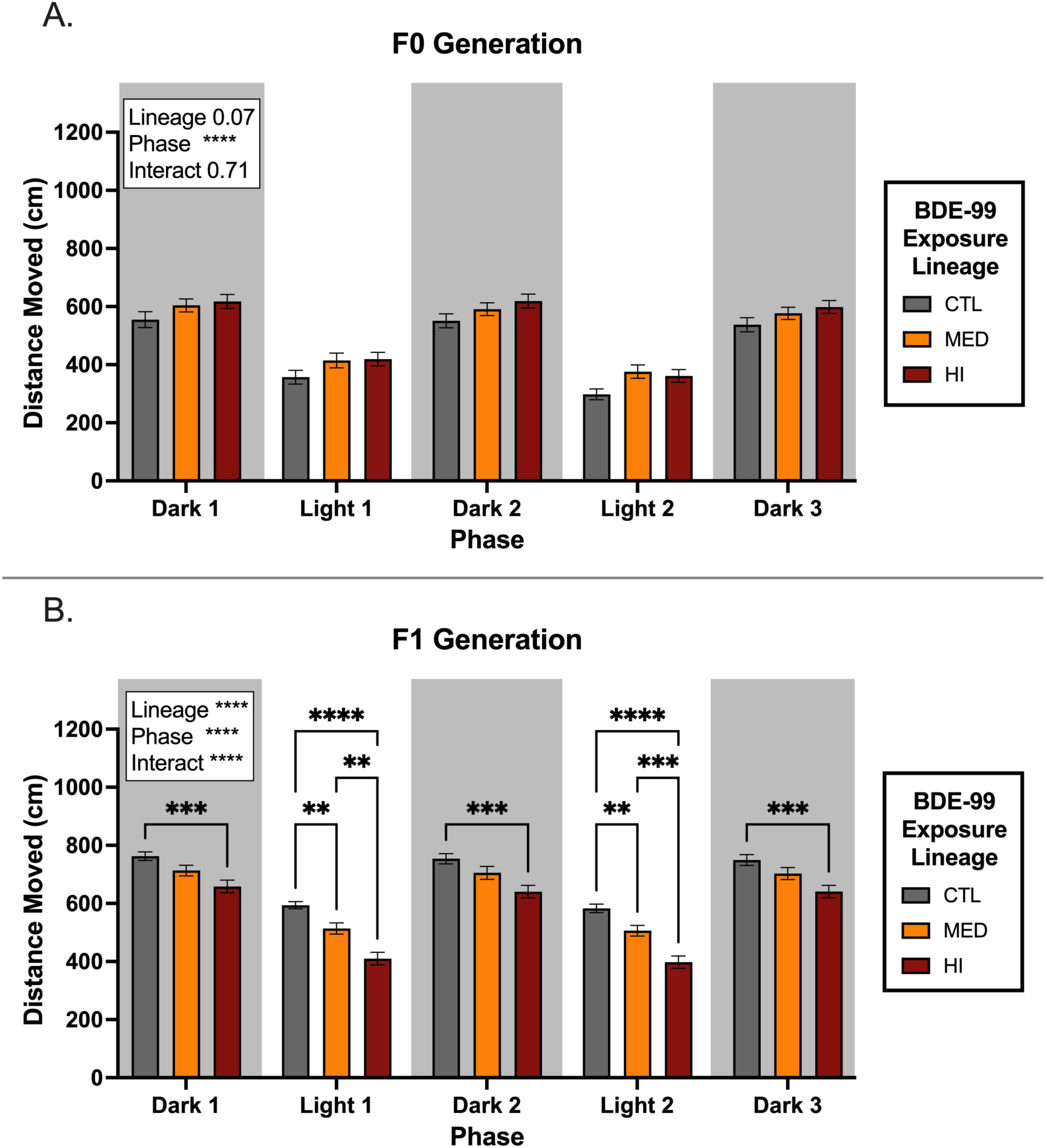
Larval photomotor responses in F0 fish that underwent direct embryonic (waterborne) exposure to BDE-99 and in their F1 descendants. Total distance moved (cm) by F0 (A) and F1 (B) larvae tracked during alternating light and dark phases. A) F0 larvae were developmentally exposed via waterborne exposure from 1-7 dpf to CTL (n=103), BDE-99 MED (n=107), or BDE-99 HI (n=102) treatments. Doses of BDE-99 in directly exposed (F0) embryos were 0 (control), 1.0, and 2.7 µg/g dry weight (dw). B) F1 larvae originated from CTL (n=57), BDE-99 MED (n=67), or BDE-99 HI (n=80) exposure lineages. Within a plot, outcomes (p-values) for statistical tests are shown for the two main effects (exposure lineage and illumination phase) and their interaction. P<0.01, p<0.001, and p<0.0001 are indicated by **, ***, and ****, respectively. Brackets with asterisks indicate significant differences between treatment means following post-hoc tests. Error bars represent standard error of the mean (SEM).

### Direct Embryonic Exposure: Anxiety-Like Behavior Effects in F0 and Their F1 Descendants

Direct embryonic exposure to BDE-99 did not affect anxiety-like behavior in juvenile F0 fish but caused subtle but statistically significant effects in F1 descendants (Figure 8). F1 descendant fish exhibited a significant main effect of exposure lineage across multiple behavioral variables, including *Total distance moved* (p=0.0351, Figure 8A). *duration spent in the top zone* (p=0.0413, Figure 8B) and *number of transitions to the top zone* (p=0.0304, Figure 8C). There was no exposure lineage effect on *Latency to enter the top zone* (Figure 8D). However, post-hoc tests indicated that treatment effects were driven by differences between the MED and HI exposure lineages, but not between exposed lineages and controls.

**Figure 8:**
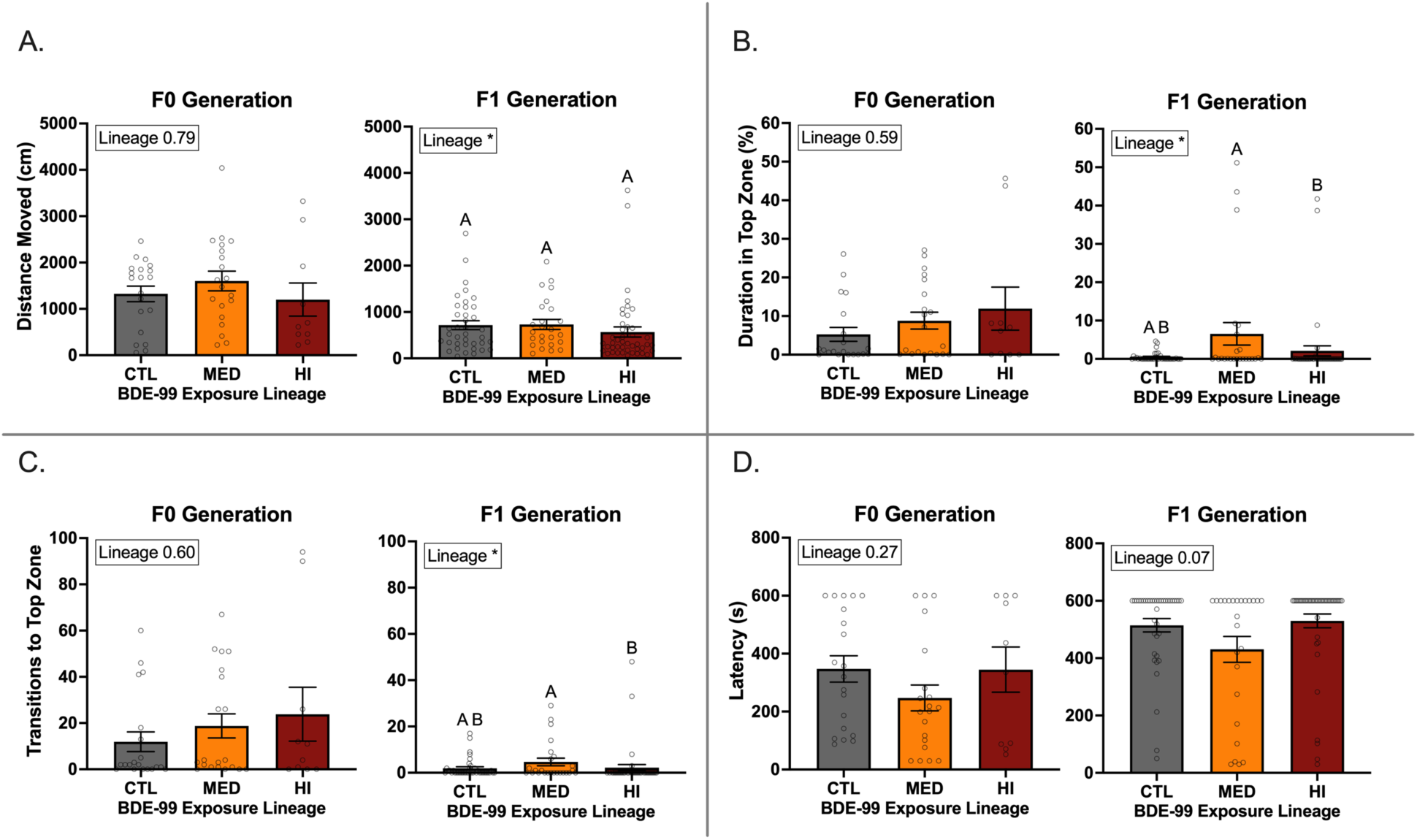
Juvenile anxiety-like behavior following direct embryonic (waterborne) exposure to BDE-99 in directly exposed (F0) and F1 descendant fish. Activity was tracked during novel tank diving tests at 4-6 months post-hatch. Variables measured include total distance moved (cm) (A), duration (% of time) spent in the top zone of the tank (B), number of transitions to the top zone (C), and latency (seconds) to enter the top zone (D). Fish that never entered the top zone were assigned the maximum latency of 600 seconds (duration of the assay). F0 fish were developmentally exposed (1-7 dpf) to CTL (n=20), BDE-99 MED (n=20), or BDE-99 HI (n=10). Doses of BDE-99 in subsets of F0 embryos were 0 (control), 1.0, and 2.7 µg/g dry weight (dw). F1 fish originated from CTL (n=38), BDE-99 MED (n=25), or BDE-99 HI (n=44) exposure lineages. Within a plot, the statistical test outcome (p-value) is shown for the main effect of exposure lineage. P<0.05 is indicated by *. The same letter above each bar indicates no significant differences between exposure lineages following post-hoc tests. Error bars represent standard error of the mean (SEM).

### Direct Embryonic Exposure: Brain Transcriptomic Responses in F1 Descendants

F1 juvenile whole-brain RNA-seq data from offspring of F0 fish that were developmentally exposed to BDE-99 were analyzed using WGCNA. Gene expression was assigned to 14 modules, each containing 60 to 283 genes. No modules were significantly associated with BDE-99 exposure lineage (adjusted p>0.05), and therefore, functional enrichment analysis was not conducted.

## Discussion

This study provides evidence that sub-lethal exposure to BDE-99 induces behavioral and molecular effects in the first-generation descendants of exposed fish, but that similar effects do not propagate into a second generation. Specifically, our findings from progenitor (F0) dietary exposure to BDE-99 demonstrate that: (1) maternal transfer of BDE-99 into eggs caused F1 descendants to be exposed starting from the earliest stages of development; (2) progenitor exposure altered both larval and juvenile behavior in F1 descendants, including photomotor and anxiety-like responses, as well as brain gene expression; (3) behavioral and molecular perturbations observed in F1 descendants were not observed in F2 descendants. Additionally, perturbations in F1 descendants occurred whether progenitors were exposed as adults (through diet) or as embryos (through water). Notably, direct embryonic exposure, which achieved comparable doses as maternal transfer, had different effects in larvae and juveniles (F0) compared to those observed in maternally-exposed F1 fish, indicating that exposure route (direct vs maternal) modulates early life neurobehavioral outcomes. This suggests that maternal effects of exposure beyond just chemical transfer are important for mediating multigenerational effects.

Dietary exposure to BDE-99 resulted in bioaccumulation over time and maternal loading of chemical into eggs, as expected for highly lipophilic chemicals. Although maternal transfer of PBDEs is well-documented in mammals^55–58,101–103^, fewer studies have examined maternal transfer of PBDEs in fish, and even fewer have quantified BDE-99 specifically. Evidence of PBDE transfer in fish exists for zebrafish^59,104^ and medaka^60^, while maternal transfer of BDE-99 specifically has been quantified in common sole^105^ and European eels^106^. However, these fish studies did not assess associated biological effects following chemical transfer, as demonstrated in the present study. We show that BDE-99 is maternally transferred in *F. heteroclitus* and correlates with behavioral outcomes in offspring. This highlights the importance of considering maternal transfer as a significant route of pollutant exposure in aquatic ecosystems. This results in exposures starting during the earliest stages of development when organisms may be particularly vulnerable to toxicological effects.

We sought to test whether the F1 effects following progenitor adult dietary exposures were entirely attributable to maternal BDE-99 transfer to developing offspring. We therefore directly exposed developing embryos (from parents that had not been exposed) to BDE-99 from their surrounding water to achieve comparable doses as those achieved through maternal transfer. However, we observed that behavioral outcomes were quite different in the two experiments. F1 fish that had been exposed through maternal transfer of BDE-99 (Experiment 1) showed extensive behavioral perturbations in larvae and juveniles, whereas F0 embryos that had been directly exposed (Experiment 2) exhibited no behavioral perturbations. Furthermore, behavioral perturbations from Experiment 2 exposures were observed in the subsequent generation, and these effects were attributable to progenitor exposure for only a short duration during early life. This suggests that the behavioral impacts of BDE-99 exposure were largely mediated through maternal inheritance of not just chemical, but also other exposure-induced changes passed from parent to offspring, and that descendant effects may emerge whether progenitors had been exposed during early or late life. Candidate mechanisms of inherited effects could include exposure-induced perturbations of energetic provisioning (yolk), maternal transfer of small molecules such as RNA, peptides, or proteins, or epigenetic imprinting. Insofar that effects did not propagate into a second generation suggests that stable exposure-induced epimutations were of limited influence, though epimutations may manifest in phenotypes that we did not measure.

The significant behavioral alterations observed in progenitor F1 fish did not follow a simple dose-response relationship. Rather, low to intermediate doses had more substantial impacts than high doses. In F1 larvae, significant hypoactivity was observed in lineages with intermediate F1 egg doses but not at the highest dose (Figure 3). Similarly, in F1 juveniles, reduced anxiety-like behaviors were detected in lineages with low to intermediate F1 egg doses but not at the highest dose. This aligns with previous research demonstrating non-monotonic behavioral effects of BDE-99. Developmental exposure of zebrafish to BDE-99 caused a non-monotonic dose effect on larval locomotor activity^66^ ^107^. Similar trends have been reported in mammals, where perinatal exposure to BDE-99 in mice altered locomotor activity at low and medium doses but not at the highest dose^108^. Non-monotonic responses to BDE-99 have also been observed in endocrine and immune effects, for example, in juvenile Chinook salmon^68,109,110^. These findings suggest that the behavioral alterations observed in our study may be part of a broader pattern of non-monotonic dose-related responses to BDE-99, consistent across multiple species and physiological systems. Understanding non-monotonic patterns like these is critical for assessing PBDE exposure risks.

We observed perturbed brain gene expression in F1 juvenile fish that were developmentally exposed to BDE-99 via maternal transfer. However, these transcriptomic changes did not consistently align with behavioral alterations. That is, behavior effects were consistently observed across most BDE-99 exposure lineage doses, yet effects on gene expression were different at different doses. Fish from medium exposure lineages displayed behavioral effects without detectable gene expression changes, while the highest concentration exhibited some gene expression shifts without corresponding behavioral alterations. One plausible explanation is that different molecular mechanisms induced in different exposure lineages may manifest behavioral perturbations that present in similar ways. Furthermore, bulk transcriptomics may be too coarse to detect molecular perturbations in specific brain regions or in minor cell types, such that single-cell RNA-seq may offer more nuanced mechanistic insight^111^. Alternatively, behavioral changes may be occurring through mechanisms independent of large-scale transcriptomic shifts, such as altered neurotransmitter signaling or synaptic plasticity. BDE-99 exposure has been associated with increased activity of the glutamate-nitric oxide-cGMP pathway^112^ and inhibition of calcium uptake in neuronal mitochondria^113^ in rat brains, as well as reduced nicotinic acetylcholine receptor (nAChR) binding in mice^114,115^, all of which can impair synaptic function. Additionally, PBDEs modulate ryanodine receptors (RyRs), further disrupting calcium homeostasis and neuronal connectivity, which may contribute to broader neurotoxic effects^116,117^. Oxidative stress may also contribute to neurobehavioral effects^118^. We speculate that the dose-dependent discrepancies in behavioral vs. gene expression changes in our study suggest multiple, perhaps redundant and interacting, molecular mechanisms may contribute to common exposure-induced neurobehavioral outcomes. Understanding the underlying molecular mechanisms is critical for defining adverse outcome pathways and assessing the full scope of BDE-99’s neurobehavioral effects.

In those experiments where we detected progenitor exposure effects on behavior, we also detected progenitor exposure effects on brain gene expression, implicating candidate causal linkages between exposure and multigenerational outcomes. Perturbed gene modules were enriched for immune responses and circadian rhythm processes. Other studies have linked PBDE exposure to immune system dysregulation. Specifically, BDE-99 exposure in Chinook salmon impaired immune function, increasing disease susceptibility and altering macrophage activity^109^. Immune system perturbations can also propagate to descendant generations through maternal transfer of antibodies and other small molecules (intergenerational immune priming). Immune system perturbations also influence behavior in fish^119^. Given the connections between immune function and neurobehavioral regulation, including modulation of neuroinflammation and stress responses, immune dysregulation may be a contributing factor to the behavioral changes observed in this study. Direct examination of impacts on immune function is warranted.

Perturbation of circadian rhythm gene expression suggests a potential link between BDE-99 exposure and circadian disruption. BDE-47 alters circadian gene expression and locomotor activity in zebrafish larvae^120^, while other structurally similar contaminants (e.g., PCB 153) also disrupt circadian-regulated gene expression in zebrafish embryos and larval behavior^121^. Since circadian regulation influences neurobehavioral processes^122–124^, these findings are consistent with the hypothesis that circadian dysregulation may contribute to BDE-99’s behavioral effects.

Our studies demonstrate neurobehavioral consequences of exposure to BDE-99, though linkages to fitness consequences could be examined with additional research. We selected neurobehavioral assays that are standard tools in biomedical and neurophysiological fields (light/dark assay probes photomotor responses, novel tank diving probes anxiety-like behavior). While the ecological implications of these effects remain uncertain, behaviors such as locomotor activity, response to stimuli, and risk avoidance are key determinants of predator-prey interactions and foraging success in natural environments^125,126^. Reduced bottom-dwelling behavior in the novel tank diving test is often interpreted as decreased anxiety-like behavior, which could indicate altered risk-avoidance strategies in natural settings^127^. Additionally, PBDE exposure has been linked to visual impairment, which could disrupt key sensory functions involved in foraging and predator detection, raising concerns about potential fitness consequences^120,128^. Since visual processing is essential for detecting environmental cues and responding to threats, even subtle disruptions could have ecological significance if they alter sensory-driven behaviors. That neurobehavioral effects from PBDE exposures manifest across generations is concerning for the health and well-being of exposed creatures, yet direct links between photomotor responses, larval hypoactivity, and anxiety-like behavior to fitness in wild populations remain limited. This presents opportunity to probe PBDE effects on other neurobehavioral endpoints, such as foraging efficiency, predator avoidance, and risk-taking tendencies^129–131^, which have stronger connections with ecological performance and fitness. Moving forward, integrating behavioral ecology paradigms and tools into toxicological research will allow for more ecologically meaningful interpretations of neurobehavioral outcomes, ultimately improving risk assessments of pollutants in aquatic systems.

Our experiments suggest limited effects that propagate to the F2. However, one caveat is that in our study, the F2 generation lineages were created with only male F1 fish from those lineages (crossed with females from unexposed control lineages), such that our experiment could only identify F2 effects of progenitor exposures if they were paternally mediated. Studies in mammals have demonstrated distinct maternal vs. paternal transmission patterns in toxicant-induced effects^132–134^. It is therefore plausible that we may have missed the detection of F2 descendant effects if they were maternally mediated.

This study provides clear evidence that BDE-99 induces intergenerational effects on behavior and brain gene expression, though these effects may not persist into a second generation. The findings contribute to a growing body of research demonstrating that maternal factors play a critical role in mediating toxicant-induced effects, likely through mechanisms beyond just chemical transfer. However, key questions remain about how parental and direct chemical exposure pathways interact, how maternal contributions shape outcomes across generations, and what additional molecular and physiological processes may be affected. Future research should aim to capture both maternal and paternal contributions to later generations, as well as examine a broader range of neurodevelopmental, behavioral, and physiological endpoints to better characterize the full multigenerational impacts of BDE-99 exposure that would matter for fitness in changing oceans. Additionally, integrating high-resolution transcriptomic approaches, such as brain-region-specific or single-cell analyses, could help identify region- or cell-type-specific mechanisms underlying observed neurobehavioral effects. Finally, these results have important implications for risk assessment of bioaccumulative pollutants in ocean ecosystems as well as in human health, emphasizing the need to consider toxicological implications of early-life exposures, maternal transfer, long-term behavioral consequences, and potential multigenerational effects.

## Data Availability

Raw RNA-seq reads are at NCBI (BioProject PRJNA1241279). A matrix of raw gene-level read counts per sample (‘bdejbrn_counts_matrix.tsv’) and RNA-seq analysis scripts are on GitHub (https://github.com/WhiteheadLab/Multigen_BDE_heteroclitus).

## Acknowledgments

Dr. Nadja Brun assisted with setting up the novel tank diving test. Jennifer Roach and Dr. Tony Gill assisted with troubleshooting the BrAD-seq library preparation method. Dr. Joanna Griffiths developed the Snakemake pipeline used to process RNA-seq data. Peyton Delaney assisted with behavioral data analysis. This research was supported by the National Institute of Environmental Health Sciences (T32 ES007059), Fumio Matsumura Memorial Endowment, Jastro-Shields Research Awards, UCD Wildlife Health Center Fellowship, Emmy Werner and Stanley Jacobsen Fellowship, Lewin Family Fellowship, and Schwall Dissertation Fellowship awarded to N.A.M.K. The views expressed in this article are those of the authors and do not necessarily represent the views or policies of the U.S. Environmental Protection Agency. Mention of trade names or commercial products does not constitute endorsement or recommendation for use.

## Supplemental Methods

We conducted two complementary experiments to investigate the multigenerational effects of exposure to BDE-99, using distinct exposure paradigms that differed in life stage and route of exposure (Figure 1). In Experiment 1, adult progenitor killifish were exposed to BDE-99 through diet, and effects were assessed across the next two generations. Experiment 2 was a complementary experiment designed to achieve similar embryonic BDE-99 doses as those measured in F1 eggs from Experiment 1. In this experiment, embryos were directly exposed to BDE-99 through waterborne exposure, allowing for a comparison of whether maternal transfer vs. direct environmental exposure influences multigenerational effects. Though many of the methods were identical between experiments, we describe them sequentially below.

### Experiment 1: Progenitor Exposure Chronological Overview

Adult wild-caught killifish were exposed to either control or BDE-99-contaminated diets for 64 days during the breeding season. Throughout this exposure period, the exposed (F0) progenitor fish were spawned to produce the parentally exposed first (F1) generation. The F1 fish were reared under uncontaminated conditions until adulthood (2 years) and subsequently spawned to produce the second (F2) generation (Figure 1A; the direct embryonic exposure experiment in Figure 1B is described later). A suite of endpoints was assessed across both generations: developmental abnormalities at 10 days post-fertilization (dpf), hatch success at 14 dpf, larval survival until 14 days post-hatch (dph), larval photomotor response (light/dark transition) at 14 dph, and juvenile anxiety-like behavior (novel tank diving) at 4-6 months post-hatch. After behavioral testing was completed, whole brain tissues from F1 and F2 fish were flash-frozen and preserved for RNA sequencing (RNA-seq).

### Adult Fish Collection and Husbandry

Adult Atlantic killifish (∼10 g wet weight (ww)) were collected from Jerusalem, Narragansett, RI, USA (41°22’58.4” N, 71°31’09.6” W) on June 6^th^, 2019 using baited traps, as previously described^1^. This site has historically served as a source of uncontaminated reference killifish, with previously measured sediment PCB concentrations ranging from 0 to 4 ng/g dry weight (dw)^2–5^. Immediately following collection, fish were transported alive to the US Environmental Protection Agency (EPA) laboratory in Narragansett. Fish were housed in large tanks (300 L, 80 gal) with a continuous flow of 23°C, 5 µm-filtered, uncontaminated seawater.

They were maintained under a 14:10 h light:dark cycle and fed TetraMin Tropical Flakes fish food (Tetra, Blacksburg, VA, USA) *ad libitum* until their use in this study one month after collection. Before the experiment, fish were marked for identification using passive integrated transponder (PIT) tags (Biomark, Boise, ID, USA) and distributed into 38-L tanks (10 gal) receiving 23°C flowing seawater. Each treatment consisted of 3 replicate tanks, with each tank containing 14 fish (8 females and 6 males). The number of individuals per tank was selected to encourage breeding to produce the next generation. Fish were held in tanks and fed an uncontaminated diet (described below) for 8 days before the initiation of the experiment. All procedures using live vertebrate animals at the EPA were conducted in accordance with Animal Care and Use Protocols approved by the EPA Institutional Animal Care and Use Committee (IACUC, ACUP # Eco23-03-002, Eco23-11-001, and Eco23-07-001).

### Diet Preparation for Progenitor BDE-99 Exposure

BDE-99 (2,2’,4,4’,5-pentabromodiphenyl ether; >99% pure) was purchased from AccuStandard (Lot no. 27669; New Haven, CT, USA). Stock solution (2.5 mg/mL) was prepared by dissolving 10 mg BDE-99 in 4 mL of acetone (Honeywell Burdick & Jackson, Muskegon, Michigan, USA). To prepare the BDE-99_MED dosing solution for one batch of diet, 52 µL of the 2.5 mg/mL BDE-99 stock was added to 1348 µL of acetone for a total volume of 1.4 mL. For the BDE-99_HI dosing solution, 209 µL of the stock was added to 1191 µL acetone for a total volume of 1.4 mL.

Each batch of diet consisted of ground TetraMin Tropical Flakes (40 g) amended with either carrier (acetone, 1.4 mL) or BDE-99 stock dissolved in acetone (1.4 mL). The 1.4 mL carrier or BDE-99 stock was added to a 1 L glass bottle, which was then rolled and allowed to evaporate before adding the ground flake food. The bottle was sealed and placed on a bottle roller for 7 days. A nutritional mixture was prepared to supplement the experimental diet. Frozen chopped spinach (454 g ww), Hikari Bio-Pure frozen brine shrimp (454 g ww), and San Francisco Bay Brand Sally’s frozen krill (454 g ww) were thawed and blended with canned chub mackerel (425 g ww) and sardines in oil (106 g ww). The mixture was divided into four portions, 450 g ww each. Each batch was supplemented with the nutritional mixture (450 g ww), solidified with gelatin (20 g dissolved in 400 mL hot DI water).

### Progenitor Dietary Exposure to BDE-99

Fish were fed acetone-amended control or contaminated diets at ∼15% of body weight for 64 days. Fish were fed treatment diets 5 days a week and control diet the other 2 days a week. Dietary treatments consisted of control and 2 concentrations of BDE-99 (37.5 and 150 ng/g fish ww/day). Target internal doses of BDE-99 were 1 and 4 μg/g fish dry weight (dw). Body burdens were measured to verify dose (see below).

### Exposed Generation Endpoints

Standard length (SL, mm) and whole fish wet weight (ww, g) were recorded for individual fish at the start (day 0) and end (day 65) of the exposure, after which Fulton’s condition index was calculated (ww/SL^3^). During this period, F0 fish were spawned at multiple time points to generate the F1 generation (spawning described below). After 64 days of feeding, all F0 (progenitor) fish were euthanized using TRICAINE-S (MS-222; Syndel, Ferndale, WA, USA), after which wet weights of gonads, liver, and abdominal fat, were measured and used to calculate the health indices gonadosomatic index (GSI), hepatosomatic index (HIS) and abdominal fat somatic index (AFSI), respectively, by dividing tissue wet weight by fish wet weight (x 100). Carcasses including tissues were divided in half (along the vertical midline).

These samples were then vacuum-sealed, flash-frozen in liquid nitrogen, and stored at -80°C for chemical analysis to quantify the body burden of BDE-99. Exposure effects on growth and health indices were tested using two-way Analysis of Variance (ANOVA) with tank as the unit of replication. The model included the main effects of BDE-99 concentration and sex and their interaction. Before ANOVA, data normality was tested (Shapiro-Wilk test). Sex effects on body burden were tested (Welch’s t-tests). All statistical analyses were conducted using GraphPad Prism version 10.4.1 for macOS (GraphPad Software, Boston, MA, USA).

### Spawning

During summer 2019 fish were manually strip-spawned 5 times following their semi-lunar spawning cycle^6^ through the exposure period to record reproductive success (yes/no) and egg counts. All eggs and milt collected from fish in a single tank were pooled to permit fertilization (F1 generation) and kept separate so that subsequent matings (to form the F2) did not include close relatives. Embryos were then incubated at 23°C under a 14:10 h light:dark cycle. Fertilization success was recorded at 4 dpf, and unfertilized eggs were archived for chemical analysis.

Fecundity, measured as the mean number of eggs per F0 female per tank at each spawning timepoint, was analyzed using a two-way repeated measures ANOVA with tank as the unit of replication. The main effects tested included BDE-99 concentration, experimental day, and their interaction. Fertilization success (percentage of fertilized F1 eggs per tank) was analyzed using a mixed-effects model with Greenhouse-Geisser correction (to account for possible sphericity violations). The model included BDE-99 concentration, experimental day, and their interaction as fixed effects. All analyses were conducted using GraphPad Prism version 10.4.1 for macOS.

In the summer of 2021, 2-year-old F1 fish were spawned to produce the F2 generation. Due to low female numbers in F1 exposure and control lineages, males were crossed with external control females. As a result, only paternal-mediated intergenerational effects were assessed in F2-generation fish. Eggs collected from 20-36 females were pooled, divided into approximately equal groups, and fertilized with pooled milt from males within each BDE-99 exposure lineage. As with the F1 generation, F2 embryos incubated at 23°C under a 14:10 h light:dark cycle. Fertilization success was recorded at 4 dpf.

### Chemical Analysis

Analytical-grade hexane and acetone was from Honeywell Burdick & Jackson. BDE-99 and ^13^C12-BDE-99 (internal standard, IS) were from Cambridge Isotope Laboratories (Tewksbury, MA, USA). Accelerated Solvent Extraction (ASE 200; Dionex Corporation, Sunnyvale, CA, USA) was used to extract BDE-99 from freeze-dried fish carcass samples and treatment diets^7^. Approximately 0.2 g of carcass was ground to a fine powder with sodium sulfate and transferred to a 22-mL stainless steel extraction cell filled with diatomaceous earth and spiked with a known amount of IS, then heated to 100°C for 5 minutes and extracted with acetone:hexane (50:50) at 1,500 psi, back-extracted with hexane, treated with sodium sulfate, and reduced in volume using ultra-high-purity nitrogen. A small portion (10%) was transferred to a pre-weighed aluminum weighing pan for lipid determination, and the remaining extract was treated with concentrated sulfuric acid.

F1 fish eggs from progenitor exposure (Experiment 1) and F0 embryos from direct embryonic exposure (Experiment 2) (40-150 mg ww) were similarly ground to a fine powder with sodium sulfate, transferred to a 2-dram scintillation vial, combined with IS and acetone, vortexed for 30 seconds, and centrifuged for 5 minutes at 2000 rpm. Extracts were back-extracted with hexane and reduced in volume. A small portion (10%) was reserved for lipid determination, and the remaining extract was treated with concentrated sulfuric acid until the hexane layer was clear.

BDE-99 concentrations in fish carcasses, eggs/embryos, and treatment diet samples were measured by performing Gas Chromatography-Mass Spectrometry (GC-MS) using an Agilent Technologies (Santa Clara, CA, USA) 6890N GC with a 5973 mass selective detector and an Agilent J&W DB-5ms Ultra Inert analytical column (15 m x 0.25 mm x 0.25 µm). Helium carrier gas was used at a constant flow rate of 1.6 mL/min. Samples were injected in splitless mode with the inlet temperature at 280°C. Ion source and quadrupole temperatures were set at 150°C and 250°C. Calibration standard solutions of BDE-99 (50-2500 ng/µL) were prepared in heptane, and a seven-point curve was generated for analysis. ChemStation software was used for data processing, and BDE-99 levels were calculated as wet, dry, and lipid weight concentrations. Procedural blanks, standard reference material (SRM 1947) analytical duplicates, and matrix spikes were routinely taken through the analytical procedure with each batch (20 samples) to evaluate and maintain quality assurance and quality control. Procedural blanks showed no contaminants. The laboratory value for BDE-99 was within 10% of the NIST-certified value. Analytical duplicates had a relative standard deviation (RSD) of <7% and matrix spike recoveries ranged from 90% to 104%.

### Embryo-Larval Rearing and Early-Life Endpoints

Embryo-Larval Assay (ELA) procedures were conducted to evaluate the developmental and survival effects of BDE-99 exposure lineage on fish embryos and larvae, following established methods with minor modifications^8,9^. Embryos were housed in clean seawater refreshed daily in Pyrex glass bowls in an incubator at 23°C with a 14:10 h light:dark cycle. At 7 dpf, embryos were transferred to individual wells in 12-well tissue culture plates (Thermo Fisher Scientific, Waltham, MA, USA) lined with clean seawater-dampened Restek Cellulose filters (20 mm, Restek, Bellefonte, PA, USA) and incubated at 23°C. At 10 dpf (late organogenesis, stage 34^10^), embryos were screened microscopically to assess the presence of developmental abnormalities, including pericardial edema, heart abnormalities, hemorrhaging in the head or tail, and reduced body or head size^11^. F1 embryos were qualitatively screened to confirm the absence of effects, while developmental abnormalities in F2 embryos were scored using a 0-7 scale (0 = no abnormalities, 7 = severe). At 14 dpf, plates were gently rocked for 1 hour to stimulate hatching, after which 3 mL of clean seawater was added to each well containing hatched larvae. Plates were checked daily to monitor late hatching and survival until 14 dph, and larvae were fed *Artemia ad libitum*. F1 and F2 embryos were treated identically, except that from 7-14 dpf, F2 embryos were maintained in wells without Restek Cellulose filters. Developmental abnormality scores in F2 embryos were analyzed using the Kruskal-Wallis test. Treatment effects on hatching success (%) and larval survival (%)were analyzed using Fisher’s exact tests for each generation.

### Behavioral Endpoints

Light/dark transition tests were conducted at 14 dph to assess larval photomotor response under alternating light and dark conditions. Assays took place between 12:00-6:00 pm. Plates containing larvae were placed on an infrared backlight (Basler AG, Ahrensburg, Germany) inside a custom-built black box, and movement was recorded from above using a Basler acA1300-60gmNIR camera. EthoVision XT software (version 11.5.1026, Noldus, Wageningen, Netherlands) was used to track activity. Each assay began with a 30-min light habituation period (100% illumination at 800 lx), followed by alternating light and dark phases: a 10-min initial dark phase (0% illumination), then two cycles of alternating 10-min phases in light (100% illumination) and dark (0% illumination). Videos were recorded at 30 frames per second and analyzed without track-smoothing applied. *Total distance moved* was calculated as cm per 10-min interval, with the initial habituation phase excluded from the analysis.

At 4-6 months post-hatch, novel tank diving tests were performed to evaluate anxiety-like behaviors in juvenile fish. Testing occurred between 12:00-6:00 pm, with counterbalanced testing times across all exposure lineages. Freshly retrieved flow-through seawater was used for all test tanks, and tank water was replaced after every six trials. The experimental setup followed a modified protocol^12^, using two adjacent rectangular tanks (33.6 cm W x 33.9 cm H x 8.2 cm D), each filled with 16.5 cm of seawater (∼4.5 L). Water temperatures were maintained at 21.5-23°C. Uniform overhead illumination (180 ± 10 lx) was provided by an LED light pad (Huion, Shenzhen, China) and verified with an LT300 light meter (EXTECH Instruments, Nashua, NH, USA). Behavioral trials were recorded using a Basler acA1300-60gmNIR camera, placed 120 cm from the front of the tanks, with video data analyzed by EthoVision XT software for fish tracking and behavioral assessment. Fish were simultaneously released into the novel tank environment and recorded for a 10-min session. The following behavioral measures were recorded: *total distance moved* (cm per 10 min), *duration spent in the top zone* of the tank (percent time), *latency to enter the top zone* (seconds), and *number of transitions to the top zone*. Subsets of juvenile F1 and F2 fish were then euthanized using MS-222, and whole brain tissues were flash-frozen in liquid nitrogen and stored at -80°C for molecular (RNA-seq) analysis.

*Total distance moved* (cm per 10 min) in the light/dark transition test was analyzed using a two-way repeated measures ANOVA, with individual larval fish as the subject (biological replicate). Greenhouse-Geisser correction was applied to account for sphericity violations. The main effects tested were BDE-99 exposure lineage, light phase, and their interaction, followed by Dunnett’s multiple comparisons tests to compare BDE-99 exposure lineages with the no-exposure control lineage. Novel tank diving measures were tested for normality (Shapiro-Wilk test). Data were analyzed using one-way ANOVA, then Tukey’s multiple comparisons test (if p<0.05) to evaluate differences across all exposure lineages. Endpoints that failed normality were analyzed using the Kruskal-Wallis test, followed by Dunn’s multiple comparisons test to compare the mean rank of each exposure lineage with that of every other lineage. Data from the F1 and F2 generations were analyzed separately. All analyses were conducted with GraphPad Prism version 10.4.1.

### Rearing to Adult

After behavioral testing at 14 dph, larvae were transferred to 9.5 L tanks (2.5 gal) at 23°C. As fish grew into juveniles, they were upgraded to 19 L (5 gal) and 38 L (10 gal) tanks to account for growth and density. Juvenile fish density was maintained at 3-10 fish per gallon in the F1 generation and 2-5 fish per gallon in the F2 generation, with the difference between generations due to variations in total fish numbers per group and space limitations.

Due to the onset of the COVID-19 pandemic, we were required to temporarily relocate F1 fish (all treatments, 6-8 months post-hatch) from the EPA facility in Narragansett, RI to the University of California, Davis for 12 months. Fish were housed in recirculating artificial seawater systems (20-22 ppt; Instant Ocean, Crystal Sea, Baltimore, MA, USA) consisting of 114 L tanks (30 gal) at a density of 0.4-0.8 fish per gallon at 20-21°C under a 12:12 h light:dark cycle. Fish were fed TetraMin Tropical Flakes food *ad libitum* in accordance with IACUC protocol #22082. In April 2021, the fish were returned to the EPA facility, where they were housed for 3 months before spawning to produce the F2 generation. Out of over 900 fish, only two died in transit to UC Davis, and one died in transit back to the EPA. Once back at the EPA facility, F1 adult fish were housed in flow-through aquaria with 23°C, 5 µm-filtered seawater at a density of 0.6-1.3 fish per gallon.

### RNA-Seq and Read Count Quantification

Juvenile fish whole brain tissues were stored at -80°C until processing. Samples were homogenized using 2.8 mm ceramic beads (Omni International, Kennesaw, GA, USA) on an Omni Bead Ruptor 24 bead mill homogenizer. The resulting lysates were suspended in 500 µL of Lysis Binding Buffer^13^, incubated at room temperature for 10 minutes, centrifuged at 13,000 rpm for 10 minutes, and the supernatant was retained. Messenger RNA (mRNA) was extracted using oligo (dT)25 beads (Dynabeads^TM^ mRNA DIRECT^TM^ Purification Kit; Invitrogen, Waltham, MA, USA) to enrich for polyadenylated transcripts. Strand-specific RNA-seq libraries were prepared using the Breath Adapter Directional sequencing (BrAD-seq) method^13^ with 13 cycles of PCR during enrichment. Fragment priming was performed with a random hexamer (GTGACTGGAGTTCAGACGTGTGCTCTTCCGATCTNNNNNNNN, where N represents a random nucleotide). Final libraries were quantified using the Qubit^TM^ dsDNA High Sensitivity Quantification Assay Kit (Invitrogen), uniquely indexed, pooled, and sequenced across six lanes of Illumina HiSeq 4000 (Illumina, San Diego, CA, USA) as paired-end 150-bp reads at the UC Davis Core Genomics Facility.

Each sample yielded approximately 9 million total raw reads. The F1 generation consisted of 4-5 biological replicates per exposure lineage, except for the 0.85 µg/g group (n = 3), for a total of 27 samples. The F2 generation consisted of 6 replicates per exposure lineage, except for the 3.50 µg/g group (n = 2), for a total of 32 samples. In total, there were six exposure lineages represented in each generation: 0, 0.85, 1.20, 2.00, and 3.50 µg/g in both generations, with 0.57 and 0.47 µg/g in F1 and F2, respectively. RNA-seq data processing was performed using a Snakemake pipeline (https://github.com/JoannaGriffiths/RNASeq-snakemake-pipeline). Raw sequencing reads were quality-checked using FastQC^14^ and trimmed with fastp^15^ to remove adapters and low-quality reads. Reads were then mapped to the *F. heteroclitus* reference genome (GenBank MU-UCD_Fhet_4.1)^16^ using Salmon^17^ for transcript-level quantification. For gene-level analyses, raw read counts were extracted and summed to the gene level using gene annotation data. RNA-seq data have been deposited in the NCBI Sequence Read Archive (BioProject: PRJNA1241279). A matrix of raw gene-level read counts per sample (‘bdejbrn_counts_matrix.tsv’) and all analysis scripts are available on GitHub (https://github.com/WhiteheadLab/Multigen_BDE_heteroclitus).

### WGCNA and Functional Enrichment

We applied weighted gene correlation network analysis (WGCNA)^18^ to identify modules of co-expressed genes that varied with experimental endpoints. This approach constructs gene networks, or modules, based on expression similarity, allowing for the detection of co-regulated gene clusters across all samples. WGCNA was conducted separately for the F1 and F2 generations to ensure independent network construction and interpretation. For each dataset, raw gene counts were normalized using variance-stabilizing transformation (VST) in DESeq2 ^19^ to account for differences in library size. Genes were then filtered based on variance, as recommended by WGNCA, retaining the top 10% most variable genes (3,164 per dataset) to reduce noise and focus on biologically relevant signals. After preprocessing, a signed adjacency matrix was constructed by calculating pairwise gene expression correlations, and an optimal soft-thresholding power was selected using the scale-free topology criterion. Gene modules, which are clusters of co-expressed genes, were identified using hierarchical clustering and dynamic tree cutting. Each module was assigned a unique color label, and eigengene values (the first principal component of module expression) were extracted to summarize module expression for each sample. Eigengene values were tested for significant associations with treatment (BDE-99 exposure lineage) using one-way ANOVA. To account for multiple comparisons across modules, p-values were adjusted using the False Discovery Rate (FDR) correction (Benjamini-Hochberg method). Modules showing significant associations (adjusted p<0.05) were further analyzed for functional enrichment of Gene Ontology (GO) terms using a Mann-Whitney *U* test^20^ to assess whether particular biological processes were overrepresented in specific gene clusters. All analyses were conducted in R (version 4.4.2)^21^ and RStudio (version 2024.09.1+394).

### Experiment 2: Direct Embryonic Exposure Chronological Overview

Fish embryos collected from unexposed parents were subjected to 6-day waterborne exposures to control or BDE-99 from 1-7 dpf (Figure 1B). These directly exposed F0 embryos were then reared to adulthood under uncontaminated conditions and subsequently spawned to produce the next (F1) generation. The same suite of developmental and behavioral endpoints tested in the progenitor exposure was assessed in both generations: developmental abnormalities at 10 dpf, hatch success at 14 dpf, larval survival until 14 dph, larval photomotor response (light/dark transition) at 14 dph, and juvenile anxiety-like behavior (novel tank diving) at 4-6 months post-hatch. Following behavioral testing, whole brains from the F1 generation were collected and archived for RNA-seq.

### Embryo Collection

Embryos were collected via mass manual spawning of lab-bred fish, originally derived (one generation previously) from a clean reference population at Scorton Creek, Sandwich, MA (41°45’53.6” N, 70°28’48.0” W)^4,22–24^. Four separate breeding tanks were used to produce four groups of embryos, which were kept separate to prevent crossing siblings or half-siblings in matings to produce the next (F1) generation. From each breeding tank, eggs were collected from 20-30 females, pooled, and fertilized with milt pooled from 20-30 males from the same tank. Several hours post-fertilization, embryos were rinsed with clean seawater and incubated at 23°C in Pyrex glass bowls. Fertilization success was assessed at 1 dpf.

### Direct Embryonic Exposure to BDE-99

A BDE-99 stock solution (2.5 mg/mL) was prepared by dissolving 10 mg of neat BDE-99 (Accustandard) in 4 mL of acetone (Honeywell Burdick & Jackson). For each exposure experiment, fertilized embryos from each of the four breeding groups were equally divided into three treatments (control and two BDE-99 concentrations) and then further divided into two replicates per treatment. As a result, each exposure experiment consisted of 24 jars (4 breeding groups x 3 treatments x 2 replicates per treatment). Embryos were mass exposed from 1 to 7 dpf in glass jars at a density of 1 embryo per 2 mL of seawater, amended with 1% dosing solution containing either carrier (acetone, control) or BDE-99 in acetone. BDE-99 concentrations were 6.2 and 26 ng/mL (labeled MED and HI, respectively). Jars were incubated at 23°C, swirled daily, and dead embryos were removed. At 7 dpf, embryos were transferred to individual wells in 12-well plates lined with clean seawater-dampened Restek Cellulose filters (20 mm) and reared as described above, in the *Embryo-Larval Rearing and Early-Life Endpoints* section. The entire exposure experiment was repeated across three separate spawning events, following the semi-lunar spawning cycle. Due to overall lower sample sizes in one of the four breeding groups, embryos from that breeding group in the third exposure experiment were archived for chemical analysis (method detailed in the *Chemical Analysis* section) to determine representative doses and uptake of BDE-99 in the embryos.

### Early-Life and Behavioral Endpoints

The same endpoints were collected for the direct embryonic exposure as the progenitor exposure and are described in detail previously in the *Embryo-Larval Rearing and Early-Life Endpoints* and *Behavioral Endpoints* sections. Endpoints included developmental abnormality scores at 10 dpf, hatch success at 14 dpf, larval photomotor response at 14 dph, larval survival until 14 dph, and juvenile anxiety-like behavior at 4-6 months post-hatch.

Developmental abnormality scores were analyzed using the Kruskal-Wallis test. Hatching success (percent hatched) and larval survival (percent alive) were analyzed using Fisher’s exact tests. *Total distance moved* (cm per 10 min) in the light/dark transition test was analyzed using a two-way repeated measures ANOVA, with individual larval fish as the subject (biological replicate) and Greenhouse-Geisser correction applied. The main effects tested were BDE-99 exposure lineage, light phase, and their interaction. Dunnett’s multiple comparisons test was performed to compare across all exposure lineages. Novel tank diving behavioral measures were tested for normality using the Shapiro-Wilk test. Endpoints were analyzed with one-way ANOVA if they passed the normality test or if they failed normality with Kruskal-Walli, followed by Dunn’s multiple comparisons test (if p<0.05) across all lineages. Data from the F0 and F1 generations were analyzed separately. All analyses were conducted with GraphPad Prism version 10.4.1.

### Rearing to Adult and Spawning

Fish from the direct embryonic exposure study were reared at the same conditions as previously described in the *Rearing to Adult* section. Juvenile fish density was maintained at 5.5-12.5 fish per gallon in the F0 generation and 1-6 fish per gallon in the F1 generation, with the difference between generations due to variations in total fish numbers per group and space limitations. As with the progenitor exposure F1 fish, the direct exposure F0 fish (all treatments) also had to be relocated to UC Davis at 6-8 months post-hatch. Rearing conditions were the same, except that density was 0.5-1 fish per gallon. Once fish were returned to the EPA facility, they were housed at a density of 0.7-1.3 fish per gallon for 3 months before spawning to produce the F1 generation.

The F1 generation was produced by crossing males and females from the different replicate groups within each exposure lineage. Therefore, effects in the F1 generation were both paternally and maternally mediated. Eggs were collected via manual strip-spawning, pooled within each replicate group, and fertilized with milt pooled from males from another replicate group in the same exposure lineage.

### RNA-Seq and WGCNA

Whole brain tissues from direct embryonic exposure F1 juvenile fish were processed as described in the *RNA-Seq and Read Count Quantification* section. Strand-specific RNA-seq libraries were prepared and sequenced together with the progenitor exposure libraries following the same method and specifications as previously described. Each sample yielded approximately 9.5 million raw reads. There were 6 replicates each from 3 breeding groups per exposure lineage, except for the MED BDE-99 lineage which had 3 and 4 replicates in 2 of the breeding groups, for a total of 49 samples. Raw sequencing reads were quality-checked and processed as previously described. WGCNA was conducted separately for the direct exposure F1 generation with all parameters and steps the same as described in the *WGCNA and Functional Enrichment* section.

---

**Figure S1:**
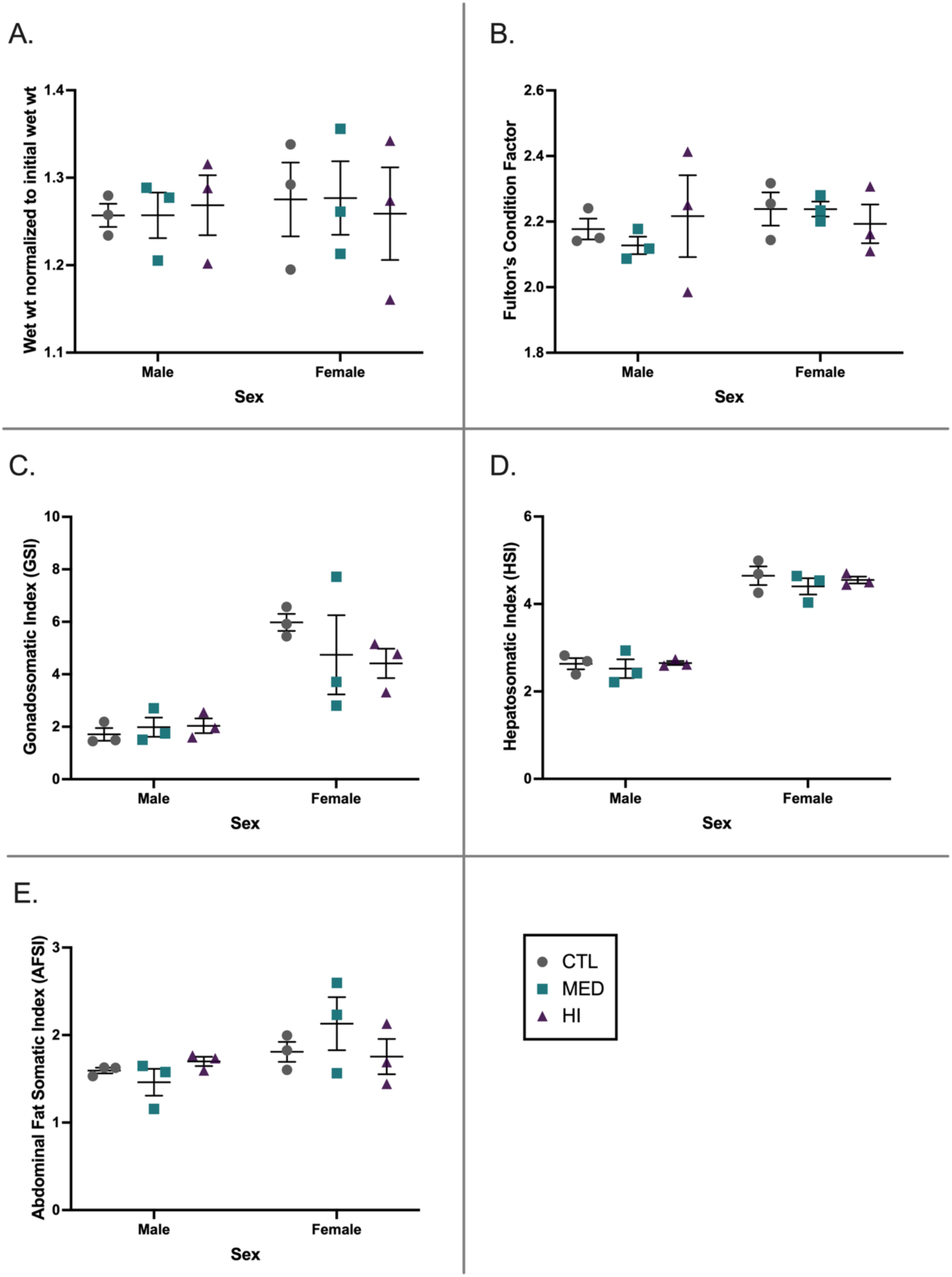
Progenitor (F0) growth and health indices by the end of the 64-day dietary exposure to BDE-99. A) Growth is expressed as the mean (± SE) of tank-level means for individual fish wet weight (ww; day 65) normalized to initial ww (day 0). B) Fulton’s condition factor was calculated as *K* = 100 x (whole fish ww [g]/standard length [mm]^3^) and is expressed as the mean (± SE) of tank-level means. C) Gonado-Somatic Index (GSI) was calculated as 100 x (gonad ww [g]/whole fish ww [g]) and is expressed as the mean (± SE) of tank-level means. Hepatosomatic Index (HSI) was calculated as 100 x (liver ww [g]/whole fish ww [g]) and is expressed as the mean (± SE) of tank-level means. Abdominal Fat Somatic Index (AFSI) was calculated as 100 x (abdominal fat ww [g]/whole fish ww [g]) and is expressed as the mean (± SE) of tank-level means. All endpoints are separated by sex and BDE-99 exposure level. Symbol colors and shapes indicate treatment groups: control (gray circles), BDE-99 MED (teal squares), and BDE-99 HI (purple triangles). Error bars represent standard error of the mean (SEM).

**Figure S2:**
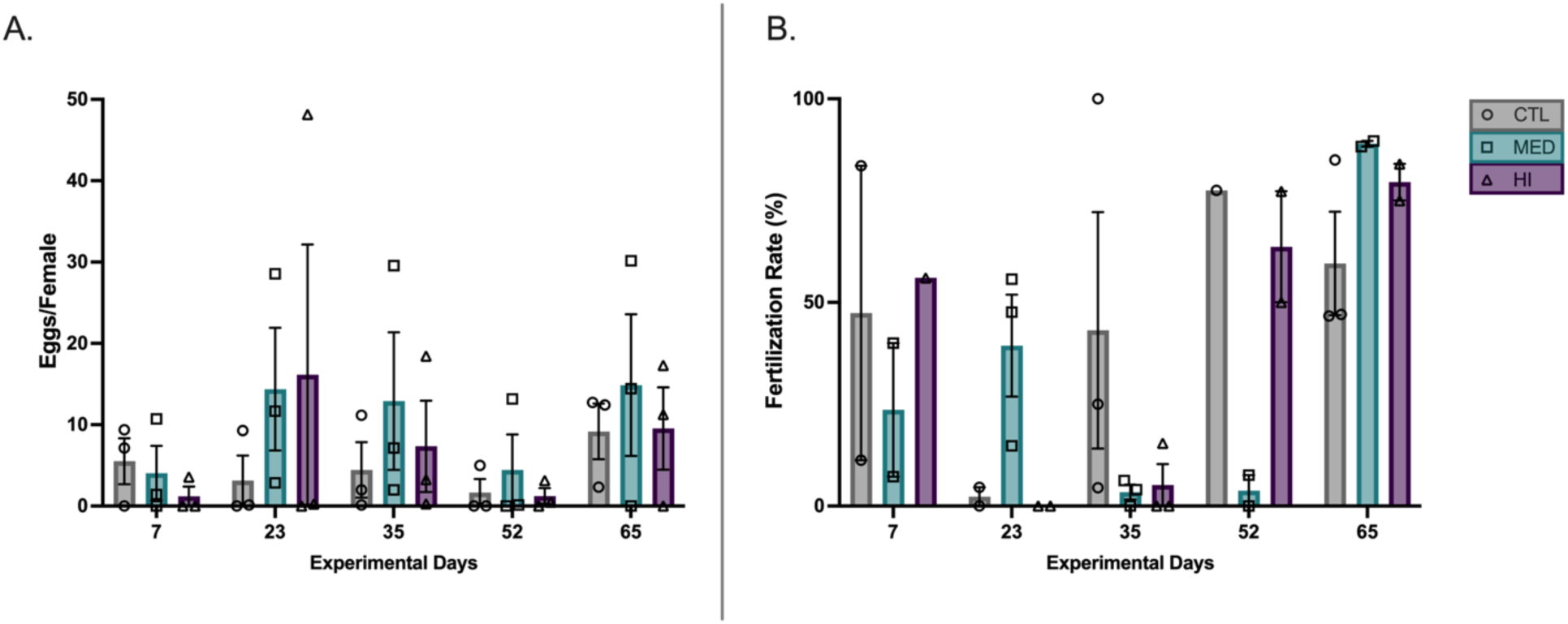
Progenitor (F0) fecundity and fertilization rate over the 64-day dietary exposure period to BDE-99. A) Fecundity was measured as number of eggs produced per female at each spawning timepoint and is expressed as the mean (± SE) of tank-level means. B) Fertilization rate was measured as the percentage of fertilized eggs produced at each spawning timepoint and is expressed as the mean (± SE) of tank-level means. Bar colors and symbol shapes indicate treatment groups: control (gray bars, circles), BDE-99 MED (teal bars, squares), and BDE-99 HI (purple bars, triangles). Error bars represent standard error of the mean (SEM).

**Figure S3:**
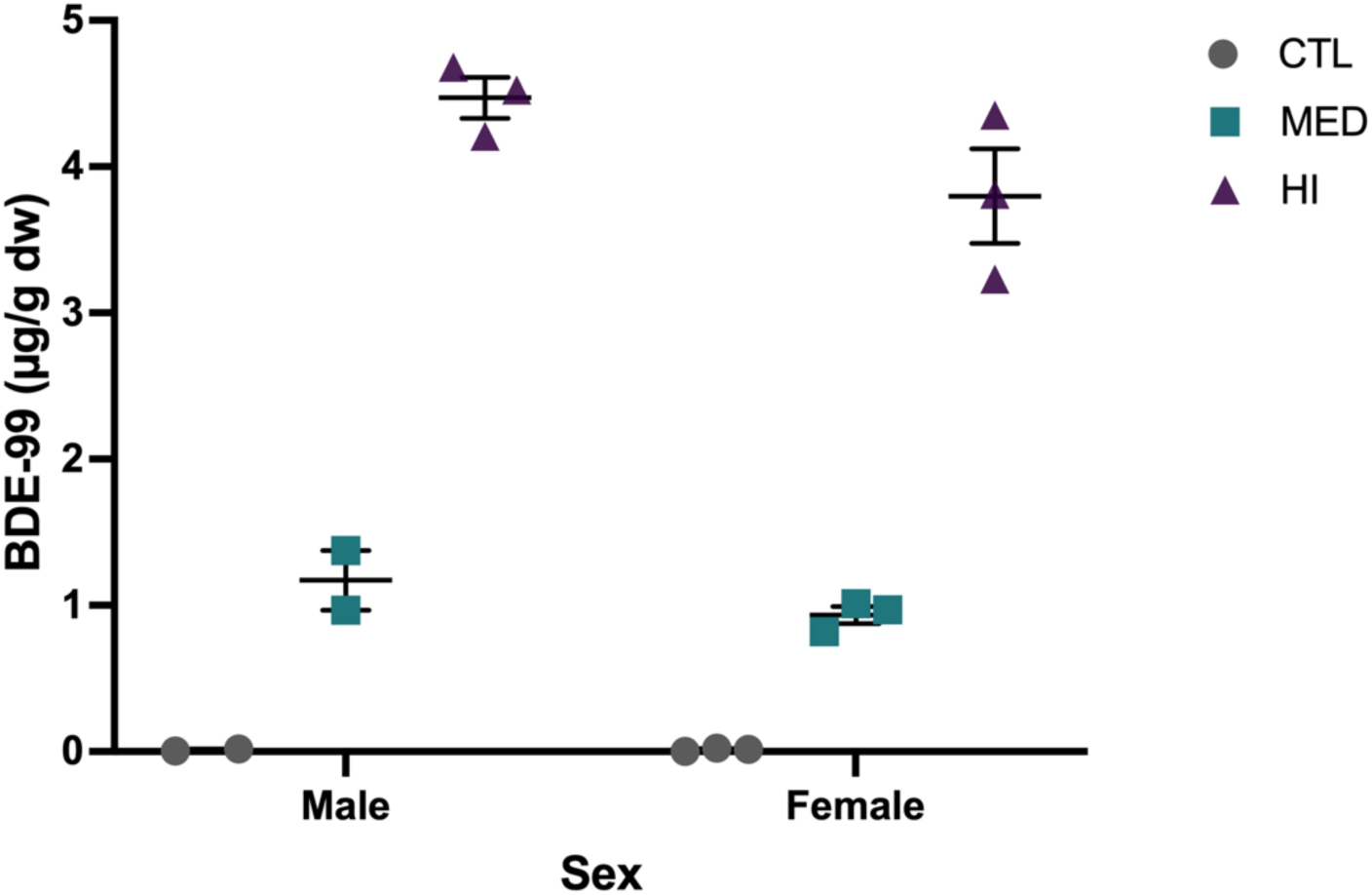
Body burdens of BDE-99 in progenitor (F0) fish by the end of the 64-day dietary exposure period. Measured body burdens of BDE-99 in fish carcasses archived on day 65 are expressed as tank-level means (µg/g dry weight [dw]), separated by sex and BDE-99 exposure level. Symbol colors and shapes indicate treatment groups: control (gray circles), BDE-99 MED (teal squares), and BDE-99 HI (purple triangles). Error bars represent standard error of the mean (SEM).

**Table S1:** Early-life endpoints in embryos and larvae from both BDE-99 exposure experiments across two generations. Developmental abnormality scores are presented as mean ± SEM, except for progenitor exposure F1, where only qualitative screening was conducted, confirming no effect. Asterisks indicate a statistically significant difference (p<0.05) between those two treatments (direct exposure F1 MED vs. HI).

| Exposure Experiment | Generation | BDE-99 Exposure Lineage | Sample Size (N) | Developmental Abnormality Score | Hatching (%) at 14 dpf | Survival (%) at 14 dph |
| --- | --- | --- | --- | --- | --- | --- |
| **Progenitor (Adult Dietary)** | **F1** | CTL | 29 | - | 89.66 | 82.76 |
| | | 0.42 $\mu\text{g/g}$ | 8 | - | 100.00 | 100.00 |
| | | 0.46 $\mu\text{g/g}$ | 19 | - | 84.21 | 84.21 |
| | | 0.47 $\mu\text{g/g}$ | 20 | - | 90.00 | 90.00 |
| | | 0.57 $\mu\text{g/g}$ | 13 | - | 100.00 | 100.00 |
| | | 2.40 $\mu\text{g/g}$ | 17 | - | 100.00 | 100.00 |
| | **F2** | CTL | 23 | 0.02 ($\pm 0.02$) | 73.91 | 86.96 |
| | | 0.47 $\mu\text{g/g}$ | 37 | 0 | 94.59 | 86.49 |
| | | 0.85 $\mu\text{g/g}$ | 69 | 0.04 ($\pm 0.02$) | 86.96 | 92.75 |
| | | 1.20 $\mu\text{g/g}$ | 25 | 0.04 ($\pm 0.05$) | 84.00 | 88.00 |
| | | 2.00 $\mu\text{g/g}$ | 43 | 0.04 ($\pm 0.06$) | 88.37 | 90.70 |
| | | 3.50 $\mu\text{g/g}$ | 23 | 0 | 73.91 | 91.30 |
| **Direct (Embryo Waterborne)** | **F0** | CTL | 166 | 0.03 ($\pm 0.02$) | 59.04 | 91.38 |
| | | MED | 171 | 0.02 ($\pm 0.02$) | 66.08 | 95.05 |
| | | HI | 166 | 0.02 ($\pm 0.01$) | 65.66 | 88.78 |
| | **F1** | CTL | 80 | 0.03 ($\pm 0.03$) | 95.00 | 78.75 |
| | | MED | 97 | 0.05 ($\pm 0.04$) | 88.66 | 89.69 * |
| | | HI | 155 | 0.09 ($\pm 0.06$) | 90.97 | 74.84 * |

## References

(1) Bell, A. M.; Hellmann, J. K. An Integrative Framework for Understanding the Mechanisms and Multigenerational Consequences of Transgenerational Plasticity. Annual Review of Ecology, Evolution, and Systematics 2019, 50 (Volume 50, 2019), 97–118. 10.1146/annurev-ecolsys-110218-024613.

(2) Turner, B. M. Epigenetic Responses to Environmental Change and Their Evolutionary Implications. Philosophical Transactions of the Royal Society B: Biological Sciences 2009, 364 (1534), 3403–3418. 10.1098/rstb.2009.0125.

(3) Wang, Y.; Liu, H.; Sun, Z. Lamarck Rises from His Grave: Parental Environment-Induced Epigenetic Inheritance in Model Organisms and Humans. Biological Reviews 2017, 92 (4), 2084–2111. 10.1111/brv.12322.

(4) Skinner, M. K. Environmental Stress and Epigenetic Transgenerational Inheritance. BMC Med 2014, 12 (1), 153. 10.1186/s12916-014-0153-y.

(5) Nilsson, E. E.; Sadler-Riggleman, I.; Skinner, M. K. Environmentally Induced Epigenetic Transgenerational Inheritance of Disease. Environmental Epigenetics 2018, 4 (2). 10.1093/eep/dvy016.

(6) Grandjean, P.; Barouki, R.; Bellinger, D. C.; Casteleyn, L.; Chadwick, L. H.; Cordier, S.; Etzel, R. A.; Gray, K. A.; Ha, E.-H.; Junien, C.; Karagas, M.; Kawamoto, T.; Lawrence, B. P.; Perera, F. P.; Prins, G. S.; Puga, A.; Rosenfeld, C. S.; Sherr, D. H.; Sly, P. D.; Suk, W.; Sun, Q.; Toppari, J.; Van Den Hazel, P.; Walker, C. L.; Heindel, J. J. Life-Long Implications of Developmental Exposure to Environmental Stressors: New Perspectives. Endocrinology 2015, 156 (10), 3408–3415. 10.1210/en.2015-1350.

(7) Anway, M. D.; Cupp, A. S.; Uzumcu, M.; Skinner, M. K. Epigenetic Transgenerational Actions of Endocrine Disruptors and Male Fertility. Science 2005, 308 (5727), 1466–1469. 10.1126/science.1108190.

(8) Anway, M. D.; Leathers, C.; Skinner, M. K. Endocrine Disruptor Vinclozolin Induced Epigenetic Transgenerational Adult-Onset Disease. Endocrinology 2006, 147 (12), 5515–5523. 10.1210/en.2006-0640.

(9) Wolstenholme, J. T.; Edwards, M.; Shetty, S. R. J.; Gatewood, J. D.; Taylor, J. A.; Rissman, E. F.; Connelly, J. J. Gestational Exposure to Bisphenol A Produces Transgenerational Changes in Behaviors and Gene Expression. Endocrinology 2012, 153 (8), 3828–3838. 10.1210/en.2012-1195.

(10) Chamorro-García, R.; Sahu, M.; Abbey, R. J.; Laude, J.; Pham, N.; Blumberg, B. Transgenerational Inheritance of Increased Fat Depot Size, Stem Cell Reprogramming, and Hepatic Steatosis Elicited by Prenatal Exposure to the Obesogen Tributyltin in Mice. Environmental Health Perspectives 2013, 121 (3), 359–366. 10.1289/ehp.1205701.

(11) Carvan III, M. J.; Kalluvila, T. A.; Klingler, R. H.; Larson, J. K.; Pickens, M.; Mora-Zamorano, F. X.; Connaughton, V. P.; Sadler-Riggleman, I.; Beck, D.; Skinner, M. K. Mercury-Induced Epigenetic Transgenerational Inheritance of Abnormal Neurobehavior Is Correlated with Sperm Epimutations in Zebrafish. PLOS ONE 2017, 12 (5), e0176155. 10.1371/journal.pone.0176155.

(12) Meyer, D. N.; Crofts, E. J.; Akemann, C.; Gurdziel, K.; Farr, R.; Baker, B. B.; Weber, D.; Baker, T. R. Developmental Exposure to Pb2+ Induces Transgenerational Changes to Zebrafish Brain Transcriptome. Chemosphere 2020, 244, 125527. 10.1016/j.chemosphere.2019.125527.

(13) Sobolewski, M.; Abston, K.; Conrad, K.; Marvin, E.; Harvey, K.; Susiarjo, M.; Cory-Slechta, D. A. Lineage- and Sex-Dependent Behavioral and Biochemical Transgenerational Consequences of Developmental Exposure to Lead, Prenatal Stress, and Combined Lead and Prenatal Stress in Mice. Environmental Health Perspectives 2020, 128 (2), 027001. 10.1289/EHP4977.

(14) Manikkam, M.; Tracey, R.; Guerrero-Bosagna, C.; Skinner, M. K. Dioxin (TCDD) Induces Epigenetic Transgenerational Inheritance of Adult Onset Disease and Sperm Epimutations. PLoS ONE 2012, 7 (9), e46249. 10.1371/journal.pone.0046249.

(15) Skinner, M. K.; Manikkam, M.; Tracey, R.; Guerrero-Bosagna, C.; Haque, M.; Nilsson, E. E. Ancestral Dichlorodiphenyltrichloroethane (DDT) Exposure Promotes Epigenetic Transgenerational Inheritance of Obesity. BMC Med 2013, 11 (1), 228. 10.1186/1741-7015-11-228.

(16) Haimbaugh, A.; Wu, C.-C.; Akemann, C.; Meyer, D. N.; Connell, M.; Abdi, M.; Khalaf, A.; Johnson, D.; Baker, T. R. Multi- and Transgenerational Effects of Developmental Exposure to Environmental Levels of PFAS and PFAS Mixture in Zebrafish (Danio Rerio). Toxics 2022, 10 (6), 334. 10.3390/toxics10060334.

(17) Baker, T. R.; Peterson, R. E.; Heideman, W. Using Zebrafish as a Model System for Studying the Transgenerational Effects of Dioxin. Toxicological Sciences 2014, 138 (2), 403–411. 10.1093/toxsci/kfu006.

(18) Knecht, A. L.; Truong, L.; Marvel, S. W.; Reif, D. M.; Garcia, A.; Lu, C.; Simonich, M. T.; Teeguarden, J. G.; Tanguay, R. L. Transgenerational Inheritance of Neurobehavioral and Physiological Deficits from Developmental Exposure to Benzo[*a*]Pyrene in Zebrafish. Toxicology and Applied Pharmacology 2017, 329, 148–157. 10.1016/j.taap.2017.05.033.

(19) Verboven, N.; Verreault, J.; Letcher, R. J.; Gabrielsen, G. W.; Evans, N. P. Differential Investment in Eggs by Arctic-Breeding Glaucous Gulls (Larus Hyperboreus) Exposed to Persistent Organic Pollutants. The Auk 2009, 126 (1), 123–133. 10.1525/auk.2009.08039.

(20) Vijendravarma, R. K.; Narasimha, S.; Kawecki, T. J. Effects of Parental Larval Diet on Egg Size and Offspring Traits in *Drosophila*. Biol. Lett. 2010, 6 (2), 238–241. 10.1098/rsbl.2009.0754.

(21) Russell, R. W.; Gobas, F. A. P. C.; Haffner, G. D. Maternal Transfer and in Ovo Exposure of Organochlorines in Oviparous Organisms: A Model and Field Verification. Environ. Sci. Technol. 1999, 33 (3), 416–420. 10.1021/es9800737.

(22) Needham, L. L.; Grandjean, P.; Heinzow, B.; Jørgensen, P. J.; Nielsen, F.; Patterson, D. G. Jr.; Sjödin, A.; Turner, W. E.; Weihe, P. Partition of Environmental Chemicals between Maternal and Fetal Blood and Tissues. Environ. Sci. Technol. 2011, 45 (3), 1121–1126. 10.1021/es1019614.

(23) Grindstaff, J. L.; Brodie, E. D.; Ketterson, E. D. Immune Function across Generations: Integrating Mechanism and Evolutionary Process in Maternal Antibody Transmission. Proc. R. Soc. Lond. B 2003, 270 (1531), 2309–2319. 10.1098/rspb.2003.2485.

(24) Groothuis, T. G. G.; Müller, W.; von Engelhardt, N.; Carere, C.; Eising, C. Maternal Hormones as a Tool to Adjust Offspring Phenotype in Avian Species. Neuroscience & Biobehavioral Reviews 2005, 29 (2), 329–352. 10.1016/j.neubiorev.2004.12.002.

(25) Hasselquist, D.; Nilsson, J.-Å. Maternal Transfer of Antibodies in Vertebrates: Trans-Generational Effects on Offspring Immunity. Phil. Trans. R. Soc. B 2009, 364 (1513), 51–60. 10.1098/rstb.2008.0137.

(26) Meylan, S.; Miles, D. B.; Clobert, J. Hormonally Mediated Maternal Effects, Individual Strategy and Global Change. Phil. Trans. R. Soc. B 2012, 367 (1596), 1647–1664. 10.1098/rstb.2012.0020.

(27) Franklin, T. B.; Russig, H.; Weiss, I. C.; Gräff, J.; Linder, N.; Michalon, A.; Vizi, S.; Mansuy, I. M. Epigenetic Transmission of the Impact of Early Stress Across Generations. Biological Psychiatry 2010, 68 (5), 408–415. 10.1016/j.biopsych.2010.05.036.

(28) Breton, C. V.; Landon, R.; Kahn, L. G.; Enlow, M. B.; Peterson, A. K.; Bastain, T.; Braun, J.; Comstock, S. S.; Duarte, C. S.; Hipwell, A.; Ji, H.; LaSalle, J. M.; Miller, R. L.; Musci, R.; Posner, J.; Schmidt, R.; Suglia, S. F.; Tung, I.; Weisenberger, D.; Zhu, Y.; Fry, R. Exploring the Evidence for Epigenetic Regulation of Environmental Influences on Child Health across Generations. Commun Biol 2021, 4 (1), 1–15. 10.1038/s42003-021-02316-6.

(29) Brander, S. M.; Biales, A. D.; Connon, R. E. The Role of Epigenomics in Aquatic Toxicology. Environmental Toxicology and Chemistry 2017, 36 (10), 2565–2573. 10.1002/etc.3930.

(30) Darnerud, P. O.; Eriksen, G. S.; Jóhannesson, T.; Larsen, P. B.; Viluksela, M. Polybrominated Diphenyl Ethers: Occurrence, Dietary Exposure, and Toxicology. Environmental Health Perspectives 2001, 109 (suppl 1), 49–68. 10.1289/ehp.01109s149.

(31) Turner, A. PBDEs in the Marine Environment: Sources, Pathways and the Role of Microplastics. Environmental Pollution 2022, 301, 118943. 10.1016/j.envpol.2022.118943.

(32) Oros, D. R.; Hoover, D.; Rodigari, F.; Crane, D.; Sericano, J. Levels and Distribution of Polybrominated Diphenyl Ethers in Water, Surface Sediments, and Bivalves from the San Francisco Estuary. Environ. Sci. Technol. 2005, 39 (1), 33–41. 10.1021/es048905q.

(33) Zhang, Y.; Wang, W.; Song, J.; Ren, Z.; Yuan, H.; Yan, H.; Zhang, J.; Pei, Z.; He, Z. Environmental Characteristics of Polybrominated Diphenyl Ethers in Marine System, with Emphasis on Marine Organisms and Sediments. BioMed Research International 2016, 2016 (1), 1317232. 10.1155/2016/1317232.

(34) Yogui, G. T.; Sericano, J. L. Polybrominated Diphenyl Ether Flame Retardants in the U.S. Marine Environment: A Review. Environment International 2009, 35 (3), 655–666. 10.1016/j.envint.2008.11.001.

(35) Rochman, C. M.; Lewison, R. L.; Eriksen, M.; Allen, H.; Cook, A.-M.; Teh, S. J. Polybrominated Diphenyl Ethers (PBDEs) in Fish Tissue May Be an Indicator of Plastic Contamination in Marine Habitats. Science of The Total Environment 2014, 476–477, 622–633. 10.1016/j.scitotenv.2014.01.058.

(36) Eljarrat, E.; Barceló, D. How Do Measured PBDE and HCBD Levels in River Fish Compare to the European Environmental Quality Standards? Environmental Research 2018, 160, 203–211. 10.1016/j.envres.2017.09.011.

(37) Li, H.; Wang, Z.; He, J.; Zhang, N.; Mao, X.; Ma, J.; Gao, H.; Yang, Z.; Ma, H. Deca-BDE Emissions, Validation, and Environmental Fate in China. Journal of Hazardous Materials 2023, 459, 132223. 10.1016/j.jhazmat.2023.132223.

(38) Muenhor, D.; Harrad, S.; Ali, N.; Covaci, A. Brominated Flame Retardants (BFRs) in Air and Dust from Electronic Waste Storage Facilities in Thailand. Environment International 2010, 36 (7), 690–698. 10.1016/j.envint.2010.05.002.

(39) Guo, J.; Lin, K.; Deng, J.; Fu, X.; Xu, Z. Polybrominated Diphenyl Ethers in Indoor Air during Waste TV Recycling Process. Journal of Hazardous Materials 2015, 283, 439–446. 10.1016/j.jhazmat.2014.09.044.

(40) Morin, N. A. O.; Andersson, P. L.; Hale, S. E.; Arp, H. P. H. The Presence and Partitioning Behavior of Flame Retardants in Waste, Leachate, and Air Particles from Norwegian Waste-Handling Facilities. Journal of Environmental Sciences 2017, 62, 115–132. 10.1016/j.jes.2017.09.005.

(41) McGrath, T. J.; Morrison, P. D.; Ball, A. S.; Clarke, B. O. Spatial Distribution of Novel and Legacy Brominated Flame Retardants in Soils Surrounding Two Australian Electronic Waste Recycling Facilities. Environ. Sci. Technol. 2018, 52 (15), 8194–8204. 10.1021/acs.est.8b02469.

(42) Hites, R. A. Polybrominated Diphenyl Ethers in the Environment and in People: A Meta-Analysis of Concentrations. Environ. Sci. Technol. 2004, 38 (4), 945–956. 10.1021/es035082g.

(43) Law, R. J.; Covaci, A.; Harrad, S.; Herzke, D.; Abdallah, M. A.-E.; Fernie, K.; Toms, L.-M. L.; Takigami, H. Levels and Trends of PBDEs and HBCDs in the Global Environment: Status at the End of 2012. Environment International 2014, 65, 147–158. 10.1016/j.envint.2014.01.006.

(44) Fromme, H.; Becher, G.; Hilger, B.; Völkel, W. Brominated Flame Retardants – Exposure and Risk Assessment for the General Population. International Journal of Hygiene and Environmental Health 2016, 219 (1), 1–23. 10.1016/j.ijheh.2015.08.004.

(45) Currier, H. A.; Fremlin, K. M.; Elliott, J. E.; Drouillard, K. G.; Williams, T. D. Bioaccumulation and Biomagnification of PBDEs in a Terrestrial Food Chain at an Urban Landfill. Chemosphere 2020, 238, 124577. 10.1016/j.chemosphere.2019.124577.

(46) Babichuk, N.; Sarkar, A.; Mulay, S.; Knight, J.; Bautista, J. J.; Young, C. J. Polybrominated Diphenyl Ethers (PBDEs) in Marine Fish and Dietary Exposure in Newfoundland. EcoHealth 2022, 19 (1), 99–113. 10.1007/s10393-022-01582-y.

(47) La Guardia, M. J.; Mainor, T. M.; Luellen, D. R.; Harvey, E.; Hale, R. C. Twenty Years Later: PBDEs in Fish from U.S. Sites with Historically Extreme Contamination. Chemosphere 2024, 351, 141126. 10.1016/j.chemosphere.2024.141126.

(48) de Wit, C. A. An Overview of Brominated Flame Retardants in the Environment. Chemosphere 2002, 46 (5), 583–624. 10.1016/S0045-6535(01)00225-9.

(49) Burreau, S.; Zebühr, Y.; Broman, D.; Ishaq, R. Biomagnification of PBDEs and PCBs in Food Webs from the Baltic Sea and the Northern Atlantic Ocean. Science of The Total Environment 2006, 366 (2), 659–672. 10.1016/j.scitotenv.2006.02.005.

(50) Mizukawa, K.; Takada, H.; Takeuchi, I.; Ikemoto, T.; Omori, K.; Tsuchiya, K. Bioconcentration and Biomagnification of Polybrominated Diphenyl Ethers (PBDEs) through Lower-Trophic-Level Coastal Marine Food Web. Marine Pollution Bulletin 2009, 58 (8), 1217–1224. 10.1016/j.marpolbul.2009.03.008.

(51) Klosterhaus, S. L.; Dreis, E.; Baker, J. E. Bioaccumulation Kinetics of Polybrominated Diphenyl Ethers from Estuarine Sediments to the Marine Polychaete, Nereis Virens. Environmental Toxicology and Chemistry 2011, 30 (5), 1204–1212. 10.1002/etc.497.

(52) Shanmuganathan, D.; Megharaj, M.; Chen, Z.; Naidu, R. Polybrominated Diphenyl Ethers (PBDEs) in Marine Foodstuffs in Australia: Residue Levels and Contamination Status of PBDEs. Marine Pollution Bulletin 2011, 63 (5), 154–159. 10.1016/j.marpolbul.2011.06.002.

(53) Domingo, J. L.; Bocio, A.; Falcó, G.; Llobet, J. M. Exposure to PBDEs and PCDEs Associated with the Consumption of Edible Marine Species. Environ. Sci. Technol. 2006, 40 (14), 4394–4399. 10.1021/es060484k.

(54) Kelly, B. C.; Ikonomou, M. G.; Blair, J. D.; Gobas, F. A. P. C. Bioaccumulation Behaviour of Polybrominated Diphenyl Ethers (PBDEs) in a Canadian Arctic Marine Food Web. Science of The Total Environment 2008, 401 (1), 60–72. 10.1016/j.scitotenv.2008.03.045.

(55) Schecter, A.; Pavuk, M.; Päpke, O.; Ryan, J. J.; Birnbaum, L.; Rosen, R. Polybrominated Diphenyl Ethers (PBDEs) in U.S. Mothers’ Milk. Environmental Health Perspectives 2003, 111 (14), 1723–1729. 10.1289/ehp.6466.

(56) Gascon, M.; Fort, M.; Martínez, D.; Carsin, A.-E.; Forns, J.; Grimalt, J. O.; Santa Marina, L.; Lertxundi, N.; Sunyer, J.; Vrijheid, M. Polybrominated Diphenyl Ethers (PBDEs) in Breast Milk and Neuropsychological Development in Infants. Environmental Health Perspectives 2012, 120 (12), 1760–1765. 10.1289/ehp.1205266.

(57) Chen, Z.-J.; Liu, H.-Y.; Cheng, Z.; Man, Y.-B.; Zhang, K.-S.; Wei, W.; Du, J.; Wong, M.-H.; Wang, H.-S. Polybrominated Diphenyl Ethers (PBDEs) in Human Samples of Mother– Newborn Pairs in South China and Their Placental Transfer Characteristics. Environment International 2014, 73, 77–84. 10.1016/j.envint.2014.07.002.

(58) Leonetti, C.; Butt, C. M.; Hoffman, K.; Miranda, M. L.; Stapleton, H. M. Concentrations of Polybrominated Diphenyl Ethers (PBDEs) and 2,4,6-Tribromophenol in Human Placental Tissues. Environment International 2016, 88, 23–29. 10.1016/j.envint.2015.12.002.

(59) Nyholm, J. R.; Norman, A.; Norrgren, L.; Haglund, P.; Andersson, P. L. Maternal Transfer of Brominated Flame Retardants in Zebrafish (*Danio Rerio*). Chemosphere 2008, 73 (2), 203–208. 10.1016/j.chemosphere.2008.04.033.

(60) van de Merwe, J. P.; Chan, A. K. Y.; Lei, E. N. Y.; Yau, M. S.; Lam, M. H. W.; Wu, R. S. S. Bioaccumulation and Maternal Transfer of PBDE 47 in the Marine Medaka (*Oryzias Melastigma*) Following Dietary Exposure. Aquatic Toxicology 2011, 103 (3), 199–204. 10.1016/j.aquatox.2011.02.021.

(61) Dietrich, J. P.; Strickland, S. A.; Hutchinson, G. P.; Van Gaest, A. L.; Krupkin, A. B.; Ylitalo, G. M.; Arkoosh, M. R. Assimilation Efficiency of PBDE Congeners in Chinook Salmon. Environ. Sci. Technol. 2015, 49 (6), 3878–3886. 10.1021/es5057038.

(62) Gómara, B.; Herrero, L.; Ramos, J. J.; Mateo, J. R.; Fernández, M. A.; García, J. F.; González, M. J. Distribution of Polybrominated Diphenyl Ethers in Human Umbilical Cord Serum, Paternal Serum, Maternal Serum, Placentas, and Breast Milk from Madrid Population, Spain. Environ. Sci. Technol. 2007, 41 (20), 6961–6968. 10.1021/es0714484.

(63) Park, J.; She, J.; Holden, A.; Sharp, M.; Gephart, R.; Souders-Mason, G.; Zhang, V.; Chow, J.; Leslie, B.; Hooper, K. High Postnatal Exposures to Polybrominated Diphenyl Ethers (PBDEs) and Polychlorinated Biphenyls (PCBs) via Breast Milk in California: Does BDE-209 Transfer to Breast Milk? Environ. Sci. Technol. 2011, 45 (10), 4579–4585. 10.1021/es103881n.

(64) Blanco, J.; Mulero, M.; Heredia, L.; Pujol, A.; Domingo, J. L.; Sánchez, D. J. Perinatal Exposure to BDE-99 Causes Learning Disorders and Decreases Serum Thyroid Hormone Levels and BDNF Gene Expression in Hippocampus in Rat Offspring. Toxicology 2013, 308, 122–128. 10.1016/j.tox.2013.03.010.

(65) Eng, M. L.; Elliott, J. E.; Williams, T. D. An Assessment of the Developmental Toxicity of BDE-99 in the European Starling Using an Integrated Laboratory and Field Approach. Ecotoxicology 2014, 23 (8), 1505–1516. 10.1007/s10646-014-1292-9.

(66) Glazer, L.; Wells, C. N.; Drastal, M.; Odamah, K.-A.; Galat, R. E.; Behl, M.; Levin, E. D. Developmental Exposure to Low Concentrations of Two Brominated Flame Retardants, BDE-47 and BDE-99, Causes Life-Long Behavioral Alterations in Zebrafish. NeuroToxicology 2018, 66, 221–232. 10.1016/j.neuro.2017.09.007.

(67) Shy, C.-G.; Huang, H.-L.; Chang-Chien, G.-P.; Chao, H.-R.; Tsou, T.-C. Neurodevelopment of Infants with Prenatal Exposure to Polybrominated Diphenyl Ethers. Bull Environ Contam Toxicol 2011, 87 (6), 643–648. 10.1007/s00128-011-0422-9.

(68) Arkoosh, M. R.; Van Gaest, A. L.; Strickland, S. A.; Hutchinson, G. P.; Krupkin, A. B.; Dietrich, J. P. Alteration of Thyroid Hormone Concentrations in Juvenile Chinook Salmon (*Oncorhynchus Tshawytscha*) Exposed to Polybrominated Diphenyl Ethers, BDE-47 and BDE-99. Chemosphere 2017, 171, 1–8. 10.1016/j.chemosphere.2016.12.035.

(69) Zhang, L.; Jin, Y.; Han, Z.; Liu, H.; Shi, L.; Hua, X.; Doering, J. A.; Tang, S.; Giesy, J. P.; Yu, H. Integrated in Silico and in Vivo Approaches to Investigate Effects of BDE-99 Mediated by the Nuclear Receptors on Developing Zebrafish. Environmental Toxicology and Chemistry 2018, 37 (3), 780–787. 10.1002/etc.4000.

(70) Berger, R. G.; Lefèvre, P. L. C.; Ernest, S. R.; Wade, M. G.; Ma, Y.-Q.; Rawn, D. F. K.; Gaertner, D. W.; Robaire, B.; Hales, B. F. Exposure to an Environmentally Relevant Mixture of Brominated Flame Retardants Affects Fetal Development in Sprague-Dawley Rats. Toxicology 2014, 320, 56–66. 10.1016/j.tox.2014.03.005.

(71) He, J.; Yang, D.; Wang, C.; Liu, W.; Liao, J.; Xu, T.; Bai, C.; Chen, J.; Lin, K.; Huang, C.; Dong, Q. Chronic Zebrafish Low Dose Decabrominated Diphenyl Ether (BDE-209) Exposure Affected Parental Gonad Development and Locomotion in F1 Offspring. Ecotoxicology 2011, 20 (8), 1813–1822. 10.1007/s10646-011-0720-3.

(72) Alfonso, S.; Blanc, M.; Joassard, L.; Keiter, S. H.; Munschy, C.; Loizeau, V.; Bégout, M.-L.; Cousin, X. Examining Multi- and Transgenerational Behavioral and Molecular Alterations Resulting from Parental Exposure to an Environmental PCB and PBDE Mixture. Aquatic Toxicology 2019, 208, 29–38. 10.1016/j.aquatox.2018.12.021.

(73) Blanc, M.; Alfonso, S.; Bégout, M.-L.; Barrachina, C.; Hyötyläinen, T.; Keiter, S. H.; Cousin, X. An Environmentally Relevant Mixture of Polychlorinated Biphenyls (PCBs) and Polybrominated Diphenylethers (PBDEs) Disrupts Mitochondrial Function, Lipid Metabolism and Neurotransmission in the Brain of Exposed Zebrafish and Their Unexposed F2 Offspring. Science of The Total Environment 2021, 754, 142097. 10.1016/j.scitotenv.2020.142097.

(74) Marteinson, S. C.; Bird, D. M.; Shutt, J. L.; Letcher, R. J.; Ritchie, I. J.; Fernie, K. J. Multi-Generational Effects of Polybrominated Diphenylethers Exposure: Embryonic Exposure of Male American Kestrels (Falco Sparverius) to DE-71 Alters Reproductive Success and Behaviors. Environmental Toxicology and Chemistry 2010, 29 (8), 1740–1747. 10.1002/etc.200.

(75) Winter, V.; Williams, T. D.; Elliott, J. E. A Three-Generational Study of In Ovo Exposure to PBDE-99 in the Zebra Finch. Environmental Toxicology and Chemistry 2013, 32 (3), 562–568. 10.1002/etc.2102.

(76) Valiela, I.; Wright, J. E.; Teal, J. M.; Volkmann, S. B. Growth, Production and Energy Transformations in the Salt-Marsh Killifish Fundulus Heteroclitus. Marine Biology 1977, 40 (2), 135–144. 10.1007/BF00396259.

(77) Nacci, D.; Coiro, L.; Champlin, D.; Jayaraman, S.; McKinney, R.; Gleason, T. R.; Munns Jr., W. R.; Specker, J. L.; Cooper, K. R. Adaptations of Wild Populations of the Estuarine Fish Fundulus Heteroclitus to Persistent Environmental Contaminants. Marine Biology 1999, 134 (1), 9–17. 10.1007/s002270050520.

(78) Rodríguez, F.; López, J. C.; Vargas, J. P.; Gómez, Y.; Broglio, C.; Salas, C. Conservation of Spatial Memory Function in the Pallial Forebrain of Reptiles and Ray-Finned Fishes. J. Neurosci. 2002, 22 (7), 2894–2903. 10.1523/JNEUROSCI.22-07-02894.2002.

(79) Rodríguez, F.; López, J. C.; Vargas, J. P.; Broglio, C.; Gómez, Y.; Salas, C. Spatial Memory and Hippocampal Pallium through Vertebrate Evolution: Insights from Reptiles and Teleost Fish. Brain Research Bulletin 2002, 57 (3), 499–503. 10.1016/S0361-9230(01)00682-7.

(80) Nacci, D. E.; Champlin, D.; Coiro, L.; McKinney, R.; Jayaraman, S. Predicting the Occurrence of Genetic Adaptation to Dioxinlike Compounds in Populations of the Estuarine Fish Fundulus Heteroclitus. Environmental Toxicology and Chemistry 2002, 21 (7), 1525–1532. 10.1002/etc.5620210726.

(81) Taylor, M. H.; Leach, G. J.; DiMichele, L.; Levitan, W. M.; Jacob, W. F. Lunar Spawning Cycle in the Mummichog, Fundulus Heteroclitus (Pisces: Cyprinodontidae). Copeia 1979, 1979 (2), 291–297. 10.2307/1443417.

(82) Stapleton, H. M.; Keller, J. M.; Schantz, M. M.; Kucklick, J. R.; Leigh, S. D.; Wise, S. A. Determination of Polybrominated Diphenyl Ethers in Environmental Standard Reference Materials. Anal Bioanal Chem 2007, 387 (7), 2365–2379. 10.1007/s00216-006-1054-5.

(83) Diane Nacci; Laura Coiro; Deena M. Wassenberg; Richard T. Di Giulio. A Non-Destructive Technique to Measure Cytochrome P4501A Enzyme Activity in Living Embryos of the Estuarine Fish Fundulus Heteroclitus. In Techniques in Aquatic Toxicology; 2005; Vol. 2.

(84) Huang, W.; Bencic, D. C.; Flick, R. L.; Nacci, D. E.; Clark, B. W.; Burkhard, L.; Lahren, T.; Biales, A. D. Characterization of the *Fundulus Heteroclitus* Embryo Transcriptional Response and Development of a Gene Expression-Based Fingerprint of Exposure for the Alternative Flame Retardant, TBPH (Bis (2-Ethylhexyl)-Tetrabromophthalate). Environmental Pollution 2019, 247, 696–705. 10.1016/j.envpol.2019.01.010.

(85) Armstrong, P. B.; Child, J. S. Stages in the Normal Development of Fundulús Heteroclitus. The Biological Bulletin 1965, 128 (2), 143–168. 10.2307/1539545.

(86) Whitehead, A.; Triant, D. A.; Champlin, D.; Nacci, D. Comparative Transcriptomics Implicates Mechanisms of Evolved Pollution Tolerance in a Killifish Population. Molecular Ecology 2010, 19 (23), 5186–5203. 10.1111/j.1365-294X.2010.04829.x.

(87) Levin, E. D.; Bencan, Z.; Cerutti, D. T. Anxiolytic Effects of Nicotine in Zebrafish. Physiology & Behavior 2007, 90 (1), 54–58. 10.1016/j.physbeh.2006.08.026.

(88) Townsley, B. T.; Covington, M. F.; Ichihashi, Y.; Zumstein, K.; Sinha, N. R. BrAD-Seq: Breath Adapter Directional Sequencing: A Streamlined, Ultra-Simple and Fast Library Preparation Protocol for Strand Specific mRNA Library Construction. Front. Plant Sci. 2015, 6. 10.3389/fpls.2015.00366.

(89) Simon Andrews. FastQC: A Quality Control tool for High Throughput Sequence Data. https://www.bioinformatics.babraham.ac.uk/projects/fastqc/ (accessed 2025-02-10).

(90) Chen, S.; Zhou, Y.; Chen, Y.; Gu, J. Fastp: An Ultra-Fast All-in-One FASTQ Preprocessor. Bioinformatics 2018, 34 (17), i884–i890. 10.1093/bioinformatics/bty560.

(91) Fundulus heteroclitus genome assembly MU-UCD_Fhet_4.1. NCBI. https://www.ncbi.nlm.nih.gov/datasets/genome/GCF_011125445.2/ (accessed 2025-02-10).

(92) Patro, R.; Duggal, G.; Love, M. I.; Irizarry, R. A.; Kingsford, C. Salmon Provides Fast and Bias-Aware Quantification of Transcript Expression. Nat Methods 2017, 14 (4), 417–419. 10.1038/nmeth.4197.

(93) Langfelder, P.; Horvath, S. WGCNA: An R Package for Weighted Correlation Network Analysis. BMC Bioinformatics 2008, 9 (1), 559. 10.1186/1471-2105-9-559.

(94) Love, M. I.; Huber, W.; Anders, S. Moderated Estimation of Fold Change and Dispersion for RNA-Seq Data with DESeq2. Genome Biology 2014, 15 (12), 550. 10.1186/s13059-014-0550-8.

(95) Wright, R. M.; Aglyamova, G. V.; Meyer, E.; Matz, M. V. Gene Expression Associated with White Syndromes in a Reef Building Coral, Acropora Hyacinthus. BMC Genomics 2015, 16 (1), 371. 10.1186/s12864-015-1540-2.

(96) R Core Team. R: A Language and Environment for Statistical Computing, 2024. https://www.R-project.org/.

(97) Susan M. Bello. Characterization of Resistance to Halogenated Aromatic Hydrocarbons in a Population of Fundulus Heteroclitus from a Marine Superfund Site. Ph.D. Thesis, Massachusetts Insitute of Technology/Woods Hole Oceanographic Institution, 1999.

(98) Bello, S. M.; Franks, D. G.; Stegeman, J. J.; Hahn, M. E. Acquired Resistance to Ah Receptor Agonists in a Population of Atlantic Killifish (Fundulus Heteroclitus) Inhabiting a Marine Superfund Site: In Vivo and in Vitro Studies on the Inducibility of Xenobiotic Metabolizing Enzymes. Toxicological Sciences 2001, 60 (1), 77–91. 10.1093/toxsci/60.1.77.

(99) Nacci, D. E.; Champlin, D.; Jayaraman, S. Adaptation of the Estuarine Fish Fundulus Heteroclitus (Atlantic Killifish) to Polychlorinated Biphenyls (PCBs). Estuaries and Coasts 2010, 33 (4), 853–864. 10.1007/s12237-009-9257-6.

(100) Fritsch, E. B.; Stegeman, J. J.; Goldstone, J. V.; Nacci, D. E.; Champlin, D.; Jayaraman, S.; Connon, R. E.; Pessah, I. N. Expression and Function of Ryanodine Receptor Related Pathways in PCB Tolerant Atlantic Killifish (*Fundulus Heteroclitus*) from New Bedford Harbor, MA, USA. Aquatic Toxicology 2015, 159, 156–166. 10.1016/j.aquatox.2014.12.017.

(101) Darnerud, P. O.; Risberg, S. Tissue Localisation of Tetra- and Pentabromodiphenyl Ether Congeners (BDE-47, -85 and -99) in Perinatal and Adult C57BL Mice. Chemosphere 2006, 62 (3), 485–493. 10.1016/j.chemosphere.2005.04.004.

(102) Frederiksen, M.; Vorkamp, K.; Mathiesen, L.; Mose, T.; Knudsen, L. E. Placental Transfer of the Polybrominated Diphenyl Ethers BDE-47, BDE-99 and BDE-209 in a Human Placenta Perfusion System: An Experimental Study. Environ Health 2010, 9 (1), 32. 10.1186/1476-069X-9-32.

(103) Koenig, C. M.; Lango, J.; Pessah, I. N.; Berman, R. F. Maternal Transfer of BDE-47 to Offspring and Neurobehavioral Development in C57BL/6J Mice. Neurotoxicology and Teratology 2012, 34 (6), 571–580. 10.1016/j.ntt.2012.09.005.

(104) Wen, Q.; Liu, H.; Zhu, Y.; Zheng, X.; Su, G.; Zhang, X.; Yu, H.; Giesy, J. P.; Lam, M. H. W. Maternal Transfer, Distribution, and Metabolism of BDE-47 and Its Related Hydroxylated, Methoxylated Analogs in Zebrafish (*Danio Rerio*). Chemosphere 2015, 120, 31–36. 10.1016/j.chemosphere.2014.05.050.

(105) Munschy, C.; Bely, N.; Héas-Moisan, K.; Olivier, N.; Loizeau, V. Tissue-Specific Distribution and Maternal Transfer of Polybrominated Diphenyl Ethers (PBDEs) and Their Metabolites in Adult Common Sole (*Solea Solea* L.) over an Entire Reproduction Cycle. Ecotoxicology and Environmental Safety 2017, 145, 457–465. 10.1016/j.ecoenv.2017.07.062.

(106) Sühring, R.; Freese, M.; Schneider, M.; Schubert, S.; Pohlmann, J.-D.; Alaee, M.; Wolschke, H.; Hanel, R.; Ebinghaus, R.; Marohn, L. Maternal Transfer of Emerging Brominated and Chlorinated Flame Retardants in European Eels. Science of The Total Environment 2015, 530–531, 209–218. 10.1016/j.scitotenv.2015.05.094.

(107) Noyes, P. D.; Haggard, D. E.; Gonnerman, G. D.; Tanguay, R. L. Advanced Morphological — Behavioral Test Platform Reveals Neurodevelopmental Defects in Embryonic Zebrafish Exposed to Comprehensive Suite of Halogenated and Organophosphate Flame Retardants. Toxicological Sciences 2015, 145 (1), 177–195. 10.1093/toxsci/kfv044.

(108) Branchi, I.; Alleva, E.; Costa, L. G. Effects of Perinatal Exposure to a Polybrominated Diphenyl Ether (PBDE 99) on Mouse Neurobehavioural Development. NeuroToxicology 2002, 23 (3), 375–384. 10.1016/S0161-813X(02)00078-5.

(109) Arkoosh, M. R.; Van Gaest, A. L.; Strickland, S. A.; Hutchinson, G. P.; Krupkin, A. B.; Dietrich, J. P. Dietary Exposure to Individual Polybrominated Diphenyl Ether Congeners BDE-47 and BDE-99 Alters Innate Immunity and Disease Susceptibility in Juvenile Chinook Salmon. Environ. Sci. Technol. 2015, 49 (11), 6974–6981. 10.1021/acs.est.5b01076.

(110) Arkoosh, M. R.; Van Gaest, A. L.; Strickland, S. A.; Hutchinson, G. P.; Krupkin, A. B.; Hicks, M. B. R.; Dietrich, J. P. Dietary Exposure to a Binary Mixture of Polybrominated Diphenyl Ethers Alters Innate Immunity and Disease Susceptibility in Juvenile Chinook Salmon (*Oncorhynchus Tshawytscha*). Ecotoxicology and Environmental Safety 2018, 163, 96–103. 10.1016/j.ecoenv.2018.07.052.

(111) Fischer, E. K.; Hauber, M. E.; Bell, A. M. Back to the Basics? Transcriptomics Offers Integrative Insights into the Role of Space, Time and the Environment for Gene Expression and Behaviour. Biology Letters 2021, 17 (9), 20210293. 10.1098/rsbl.2021.0293.

(112) Llansola, M.; Erceg, S.; Monfort, P.; Montoliu, C.; Felipo, V. Prenatal Exposure to Polybrominated Diphenylether 99 Enhances the Function of the Glutamate–Nitric Oxide– cGMP Pathway in Brain in Vivo and in Cultured Neurons. European Journal of Neuroscience 2007, 25 (2), 373–379. 10.1111/j.1460-9568.2006.05289.x.

(113) Coburn, C. G.; Currás-Collazo, M. C.; Kodavanti, P. R. S. In Vitro Effects of Environmentally Relevant Polybrominated Diphenyl Ether (PBDE) Congeners on Calcium Buffering Mechanisms in Rat Brain. Neurochem Res 2008, 33 (2), 355–364. 10.1007/s11064-007-9430-x.

(114) Viberg, H.; Fredriksson, A.; Eriksson, P. Neonatal Exposure to the Brominated Flame Retardant 2,2’,4,4’,5-Pentabromodiphenyl Ether Causes Altered Susceptibility in the Cholinergic Transmitter System in the Adult Mouse. Toxicological Sciences 2002, 67 (1), 104–107. 10.1093/toxsci/67.1.104.

(115) Viberg, H.; Fredriksson, A.; Eriksson, P. Neonatal Exposure to the Brominated Flame-Retardant, 2,2′,4,4′,5-Pentabromodiphenyl Ether, Decreases Cholinergic Nicotinic Receptors in Hippocampus and Affects Spontaneous Behaviour in the Adult Mouse. Environmental Toxicology and Pharmacology 2004, 17 (2), 61–65. 10.1016/j.etap.2004.02.004.

(116) Kim, K. H.; Bose, D. D.; Ghogha, A.; Riehl, J.; Zhang, R.; Barnhart, C. D.; Lein, P. J.; Pessah, I. N. Para- and Ortho-Substitutions Are Key Determinants of Polybrominated Diphenyl Ether Activity toward Ryanodine Receptors and Neurotoxicity. Environmental Health Perspectives 2011, 119 (4), 519–526. 10.1289/ehp.1002728.

(117) Chen, H.; Streifel, K. M.; Singh, V.; Yang, D.; Mangini, L.; Wulff, H.; Lein, P. J. From the Cover: BDE-47 and BDE-49 Inhibit Axonal Growth in Primary Rat Hippocampal Neuron-Glia Co-Cultures via Ryanodine Receptor-Dependent Mechanisms. Toxicological Sciences 2017, 156 (2), 375–386. 10.1093/toxsci/kfw259.

(118) Cheng, J.; Gu, J.; Ma, J.; Chen, X.; Zhang, M.; Wang, W. Neurobehavioural Effects, Redox Responses and Tissue Distribution in Rat Offspring Developmental Exposure to BDE-99. Chemosphere 2009, 75 (7), 963–968. 10.1016/j.chemosphere.2009.01.004.

(119) Kirsten, K.; Fior, D.; Kreutz, L. C.; Barcellos, L. J. G. First Description of Behavior and Immune System Relationship in Fish. Sci Rep 2018, 8 (1), 846. 10.1038/s41598-018-19276-3.

(120) Cao, M.; Xu, T.; Zhang, H.; Wei, S.; Wang, H.; Song, Y.; Guo, X.; Chen, D.; Yin, D. BDE-47 Causes Depression-like Effects in Zebrafish Larvae via a Non-Image-Forming Visual Mechanism. Environ. Sci. Technol. 2023, 57 (26), 9592–9602. 10.1021/acs.est.3c01716.

(121) Aluru, N.; Krick, K. S.; McDonald, A. M.; Karchner, S. I. Developmental Exposure to PCB153 (2,2’,4,4’,5,5’-Hexachlorobiphenyl) Alters Circadian Rhythms and the Expression of Clock and Metabolic Genes. Toxicological Sciences 2020, 173 (1), 41–52. 10.1093/toxsci/kfz217.

(122) Easton, A.; Arbuzova, J.; Turek, F. W. The Circadian Clock Mutation Increases Exploratory Activity and Escape-Seeking Behavior. *Genes*, Brain and Behavior 2003, 2 (1), 11–19. 10.1034/j.1601-183X.2003.00002.x.

(123) Roybal, K.; Theobold, D.; Graham, A.; DiNieri, J. A.; Russo, S. J.; Krishnan, V.; Chakravarty, S.; Peevey, J.; Oehrlein, N.; Birnbaum, S.; Vitaterna, M. H.; Orsulak, P.; Takahashi, J. S.; Nestler, E. J.; Carlezon, W. A.; McClung, C. A. Mania-like Behavior Induced by Disruption of CLOCK. Proceedings of the National Academy of Sciences 2007, 104 (15), 6406–6411. 10.1073/pnas.0609625104.

(124) Barnard, A. R.; Nolan, P. M. When Clocks Go Bad: Neurobehavioural Consequences of Disrupted Circadian Timing. PLOS Genetics 2008, 4 (5), e1000040. 10.1371/journal.pgen.1000040.

(125) Scott, G. R.; Sloman, K. A. The Effects of Environmental Pollutants on Complex Fish Behaviour: Integrating Behavioural and Physiological Indicators of Toxicity. Aquatic Toxicology 2004, 68 (4), 369–392. 10.1016/j.aquatox.2004.03.016.

(126) Jacquin, L.; Petitjean, Q.; Côte, J.; Laffaille, P.; Jean, S. Effects of Pollution on Fish Behavior, Personality, and Cognition: Some Research Perspectives. Front. Ecol. Evol. 2020, 8, 86. 10.3389/fevo.2020.00086.

(127) Vorhees, C. V.; Williams, M. T.; Hawkey, A. B.; Levin, E. D. Translating Neurobehavioral Toxicity Across Species From Zebrafish to Rats to Humans: Implications for Risk Assessment. Front. Toxicol. 2021, 3. 10.3389/ftox.2021.629229.

(128) Wei, S.; Chen, F.; Xu, T.; Cao, M.; Yang, X.; Zhang, B.; Guo, X.; Yin, D. BDE-99 Disrupts the Photoreceptor Patterning of Zebrafish Larvae via Transcription Factor Six7. Environ. Sci. Technol. 2022, 56 (9), 5673–5683. 10.1021/acs.est.1c08914.

(129) Rice, D.; Barone, S. Critical Periods of Vulnerability for the Developing Nervous System: Evidence from Humans and Animal Models. Environmental Health Perspectives 2000, 108 (suppl 3), 511–533. 10.1289/ehp.00108s3511.

(130) Costa, L. G.; Giordano, G. Developmental Neurotoxicity of Polybrominated Diphenyl Ether (PBDE) Flame Retardants. NeuroToxicology 2007, 28 (6), 1047–1067. 10.1016/j.neuro.2007.08.007.

(131) Grandjean, P.; Landrigan, P. J. Neurobehavioural Effects of Developmental Toxicity. The Lancet Neurology 2014, 13 (3), 330–338. 10.1016/S1474-4422(13)70278-3.

(132) Magnus, M. C.; Håberg, S. E.; Karlstad, Ø.; Nafstad, P.; London, S. J.; Nystad, W. Grandmother’s Smoking When Pregnant with the Mother and Asthma in the Grandchild: The Norwegian Mother and Child Cohort Study. 2015. 10.1136/thoraxjnl-2014-206438.

(133) Accordini, S.; Calciano, L.; Johannessen, A.; Portas, L.; Benediktsdóttir, B.; Bertelsen, R. J.; Bråbäck, L.; Carsin, A.-E.; Dharmage, S. C.; Dratva, J.; Forsberg, B.; Gomez Real, F.; Heinrich, J.; Holloway, J. W.; Holm, M.; Janson, C.; Jögi, R.; Leynaert, B.; Malinovschi, A.; Marcon, A.; Martínez-Moratalla Rovira, J.; Raherison, C.; Sánchez-Ramos, J. L.; Schlünssen, V.; Bono, R.; Corsico, A. G.; Demoly, P.; Dorado Arenas, S.; Nowak, D.; Pin, I.; Weyler, J.; Jarvis, D.; Svanes, C.; the Ageing Lungs in European Cohorts (ALEC) Study. A Three-Generation Study on the Association of Tobacco Smoking with Asthma. International Journal of Epidemiology 2018, 47 (4), 1106–1117. 10.1093/ije/dyy031.

(134) Wolstenholme, J. T.; Drobná, Z.; Henriksen, A. D.; Goldsby, J. A.; Stevenson, R.; Irvin, J. W.; Flaws, J. A.; Rissman, E. F. Transgenerational Bisphenol A Causes Deficits in Social Recognition and Alters Postsynaptic Density Genes in Mice. Endocrinology 2019, 160 (8), 1854–1867. 10.1210/en.2019-00196.

## References (for Supplemental Methods)

(1) Nacci, D. E.; Champlin, D.; Coiro, L.; McKinney, R.; Jayaraman, S. Predicting the Occurrence of Genetic Adaptation to Dioxinlike Compounds in Populations of the Estuarine Fish Fundulus Heteroclitus. Environmental Toxicology and Chemistry 2002, 21 (7), 1525–1532. 10.1002/etc.5620210726.

(2) McMillan, A. M.; Bagley, M. J.; Jackson, S. A.; Nacci, D. E. Genetic Diversity and Structure of an Estuarine Fish (Fundulus Heteroclitus) Indigenous to Sites Associated with a Highly Contaminated Urban Harbor. Ecotoxicology 2006, 15 (6), 539–548. 10.1007/s10646-006-0090-4.

(3) Bozinovic, G.; Oleksiak, M. F. Embryonic Gene Expression among Pollutant Resistant and Sensitive *Fundulus Heteroclitus* Populations. Aquatic Toxicology 2010, 98 (3), 221–229. 10.1016/j.aquatox.2010.02.022.

(4) Nacci, D. E.; Champlin, D.; Jayaraman, S. Adaptation of the Estuarine Fish Fundulus Heteroclitus (Atlantic Killifish) to Polychlorinated Biphenyls (PCBs). Estuaries and Coasts 2010, 33 (4), 853–864. 10.1007/s12237-009-9257-6.

(5) Markert, J. A.; Rock, M. T.; Clark, B. W.; Nacci, D. E. Urbanization and Distance Shape Population Structure in Fundulus Heteroclitus. Journal of Urban Ecology 2024, 10 (1), juae016. 10.1093/jue/juae016.

(6) Taylor, M. H.; Leach, G. J.; DiMichele, L.; Levitan, W. M.; Jacob, W. F. Lunar Spawning Cycle in the Mummichog, Fundulus Heteroclitus (Pisces: Cyprinodontidae). Copeia 1979, 1979 (2), 291–297. 10.2307/1443417.

(7) Stapleton, H. M.; Keller, J. M.; Schantz, M. M.; Kucklick, J. R.; Leigh, S. D.; Wise, S. A. Determination of Polybrominated Diphenyl Ethers in Environmental Standard Reference Materials. Anal Bioanal Chem 2007, 387 (7), 2365–2379. 10.1007/s00216-006-1054-5.

(8) Diane Nacci; Laura Coiro; Deena M. Wassenberg; Richard T. Di Giulio. A Non-Destructive Technique to Measure Cytochrome P4501A Enzyme Activity in Living Embryos of the Estuarine Fish Fundulus Heteroclitus. In Techniques in Aquatic Toxicology; 2005; Vol. 2.

(9) Huang, W.; Bencic, D. C.; Flick, R. L.; Nacci, D. E.; Clark, B. W.; Burkhard, L.; Lahren, T.; Biales, A. D. Characterization of the *Fundulus Heteroclitus* Embryo Transcriptional Response and Development of a Gene Expression-Based Fingerprint of Exposure for the Alternative Flame Retardant, TBPH (Bis (2-Ethylhexyl)-Tetrabromophthalate). Environmental Pollution 2019, 247, 696–705. 10.1016/j.envpol.2019.01.010.

(10) Armstrong, P. B.; Child, J. S. Stages in the Normal Development of Fundulús Heteroclitus. The Biological Bulletin 1965, 128 (2), 143–168. 10.2307/1539545.

(11) Whitehead, A.; Triant, D. A.; Champlin, D.; Nacci, D. Comparative Transcriptomics Implicates Mechanisms of Evolved Pollution Tolerance in a Killifish Population. Molecular Ecology 2010, 19 (23), 5186–5203. 10.1111/j.1365-294X.2010.04829.x.

(12) Levin, E. D.; Bencan, Z.; Cerutti, D. T. Anxiolytic Effects of Nicotine in Zebrafish. Physiology & Behavior 2007, 90 (1), 54–58. 10.1016/j.physbeh.2006.08.026.

(13) Townsley, B. T.; Covington, M. F.; Ichihashi, Y.; Zumstein, K.; Sinha, N. R. BrAD-Seq: Breath Adapter Directional Sequencing: A Streamlined, Ultra-Simple and Fast Library Preparation Protocol for Strand Specific mRNA Library Construction. Front. Plant Sci. 2015, 6. 10.3389/fpls.2015.00366.

(14) Simon Andrews. FastQC: A Quality Control tool for High Throughput Sequence Data. https://www.bioinformatics.babraham.ac.uk/projects/fastqc/ (accessed 2025-02-10).

(15) Chen, S.; Zhou, Y.; Chen, Y.; Gu, J. Fastp: An Ultra-Fast All-in-One FASTQ Preprocessor. Bioinformatics 2018, 34 (17), i884–i890. 10.1093/bioinformatics/bty560.

(16) Fundulus heteroclitus genome assembly MU-UCD_Fhet_4.1. NCBI. https://www.ncbi.nlm.nih.gov/datasets/genome/GCF_011125445.2/ (accessed 2025-02-10).

(17) Patro, R.; Duggal, G.; Love, M. I.; Irizarry, R. A.; Kingsford, C. Salmon Provides Fast and Bias-Aware Quantification of Transcript Expression. Nat Methods 2017, 14 (4), 417–419. 10.1038/nmeth.4197.

(18) Langfelder, P.; Horvath, S. WGCNA: An R Package for Weighted Correlation Network Analysis. BMC Bioinformatics 2008, 9 (1), 559. 10.1186/1471-2105-9-559.

(19) Love, M. I.; Huber, W.; Anders, S. Moderated Estimation of Fold Change and Dispersion for RNA-Seq Data with DESeq2. Genome Biology 2014, 15 (12), 550. 10.1186/s13059-014-0550-8.

(20) Wright, R. M.; Aglyamova, G. V.; Meyer, E.; Matz, M. V. Gene Expression Associated with White Syndromes in a Reef Building Coral, Acropora Hyacinthus. BMC Genomics 2015, 16 (1), 371. 10.1186/s12864-015-1540-2.

(21) R Core Team. R: A Language and Environment for Statistical Computing, 2024. https://www.R-project.org/.

(22) Susan M. Bello. Characterization of Resistance to Halogenated Aromatic Hydrocarbons in a Population of Fundulus Heteroclitus from a Marine Superfund Site. Ph.D. Thesis, Massachusetts Insitute of Technology/Woods Hole Oceanographic Institution, 1999.

(23) Bello, S. M.; Franks, D. G.; Stegeman, J. J.; Hahn, M. E. Acquired Resistance to Ah Receptor Agonists in a Population of Atlantic Killifish (Fundulus Heteroclitus) Inhabiting a Marine Superfund Site: In Vivo and in Vitro Studies on the Inducibility of Xenobiotic Metabolizing Enzymes. Toxicological Sciences 2001, 60 (1), 77–91. 10.1093/toxsci/60.1.77.

(24) Fritsch, E. B.; Stegeman, J. J.; Goldstone, J. V.; Nacci, D. E.; Champlin, D.; Jayaraman, S.; Connon, R. E.; Pessah, I. N. Expression and Function of Ryanodine Receptor Related Pathways in PCB Tolerant Atlantic Killifish (*Fundulus Heteroclitus*) from New Bedford Harbor, MA, USA. Aquatic Toxicology 2015, 159, 156–166. 10.1016/j.aquatox.2014.12.017.

